# Single-molecule analysis of receptor-β-arrestin interactions in living cells

**DOI:** 10.1101/2022.11.15.516577

**Authors:** Jak Grimes, Zsombor Koszegi, Yann Lanoiselée, Tamara Miljus, Shannon L. O’Brien, Tomasz M Stepniewski, Brian Medel-Lacruz, Mithu Baidya, Maria Makarova, Dylan M. Owen, Arun K. Shukla, Jana Selent, Stephen J. Hill, Davide Calebiro

## Abstract

β-arrestin plays a key role in G protein-coupled receptor (GPCR) signaling and desensitization. Despite recent structural advances, the mechanisms that govern receptor–β-arrestin interactions at the plasma membrane of living cells remain elusive. Here, we combine single-molecule microscopy with molecular dynamics simulations to dissect the complex sequence of events involved in β-arrestin interactions with both receptors and the lipid bilayer. In contrast to the currently widely accepted model, we show that β-arrestin spontaneously inserts into the lipid bilayer and transiently interacts with receptors via lateral diffusion on the plasma membrane. Moreover, we show that following receptor interaction, the plasma membrane stabilizes β-arrestin in a membrane-bound, active-like conformation, allowing it to diffuse to clathrin coated pits separately from the activating receptor. These results challenge our current understanding of β-arrestin function at the plasma membrane, revealing a new critical role for β-arrestin pre-association with the lipid bilayer in facilitating its interactions with receptors and subsequent activation.

## INTRODUCTION

G protein coupled receptors (GPCRs), the largest family of cell receptors, are implicated in virtually every physiological process and are major drug targets (Hauser et al., 2017; Pierce et al., 2002). GPCR signaling and its regulation largely rely upon interactions of the receptors embedded in the plasma membrane with two classes of signaling molecules, i.e. G proteins and β-arrestins (Pierce et al., 2002).

Following agonist-mediated GPCR activation and phosphorylation by G protein coupled-receptor kinases (GRKs), β-arrestins located in the cytoplasm are thought to translocate and bind directly to the agonist occupied, phosphorylated receptors on the plasma membrane. By interacting with the receptor core, β-arrestins interfere with G protein coupling, thus mediating rapid signal desensitization (Lohse et al., 1990). In addition, β-arrestins trigger receptor internalization via interaction with the adaptor protein 2 (AP2) and clathrin heavy chain (Pierce and Lefkowitz, 2001). Besides these ‘classical’ functions, β-arrestins have been proposed to mediate ‘non-classical’ G protein-independent effects (Daaka et al., 1998), providing a mechanism for ‘biased’ signaling (Kumari et al., 2016; Reiter et al., 2012). Moreover, recent findings of prolonged β-arrestin activation (Eichel et al., 2018; Latorraca et al., 2018; Nuber et al., 2016) suggest that β-arrestin signaling might be more complex than previously thought.

Although recent studies with purified proteins have provided new important structural insights into receptor–arrestin interactions (Huang et al., 2020; Kang et al., 2015; Lee et al., 2020; Staus et al., 2020; Thomsen et al., 2016; Zhou et al., 2017), the complex and, likely, highly dynamic sequence of events involved in β-arrestin interaction with GPCRs on the plasma membrane of living cells remains largely unknown. This includes fundamental and presently unanswered questions such as: How do GPCRs and β-arrestins meet on the plasma membrane? How long do they interact? What are the molecular mechanisms underlying prolonged β-arrestin activation? How do β-arrestins and receptors reach clathrin coated pits (CCPs) to mediate GPCR internalization and ‘non-classical’ signaling?

Despite considerable efforts, answering these questions has been hampered by important limitations of the conventional approaches commonly used to investigate GPCR signaling, which include biochemical and ensemble biophysical methods. A first limitation is their insufficient spatio-temporal resolution, which precludes a detailed investigation of protein– protein interactions on the plasma membrane as these typically occur on scales of tens of nanometers and milliseconds (Calebiro and Sungkaworn, 2017). A second important limitation is that ensemble methods only report the average behavior of thousands or millions of generally unsynchronized molecules, precluding a direct estimation of kinetics parameters or the analysis of complex sequences of events (Calebiro and Sungkaworn, 2017).

To circumvent these limitations, here we combined an innovative multicolor single-molecule microscopy approach in living cells – which we previously used to investigate receptor–G protein interactions (Sungkaworn et al., 2017) – with molecular dynamics (MD) simulations and other advanced computational approaches to dissect the precise sequence of events involved in receptor–β-arrestin interactions. Importantly, this allowed us to directly visualize and investigate individual β-arrestins and receptors as they diffuse and interact at the plasma membrane of living cells with unprecedented spatial (~20 nm) and temporal (~30 ms) resolution. As a first model, we chose β-arrestin 2 (βArr2) and the β_2_-adrenergic receptor (β_2_AR), a prototypical family-A GPCR that controls numerous physiological processes, including regulation of smooth muscle tone and heart contractility (Lohse et al., 2003; Zhang and Mende, 2011). Furthermore, we investigated βArr2 with two additional receptors, the β_1_-adrenergic receptor (β_1_AR) and a chimeric β_2_AR–vasopressin V_2_ receptor (β_2_V_2_), which have been extensively used as models of weak and strong β-arrestin interactions, respectively (Oakley et al., 1999; Shukla et al., 2014).

Unexpectedly, our results show that, contrary to the currently widely accepted model, β-arrestin does not directly translocate to receptors from the cytoplasm but rather spontaneously inserts into the plasma membrane via hydrophobic interactions mediated by its C-edge and that this pre-association is critical for β-arrestin to engage with receptors via lateral diffusion. Moreover, they reveal that receptor–β-arrestin interactions are much more dynamic than previously thought, lasting on average only approximately 1 s, a time insufficient for receptors and β-arrestins to reach CCPs together as currently assumed. Finally, they uncover a previously unknown potential interaction of the finger loop domain of β-arrestin with the plasma membrane, which appears to play a critical role in stabilizing β-arrestin in a membrane-bound, active-like conformation after transient interaction with a receptor.

## RESULTS

### Single-molecule imaging reveals spontaneous membrane translocation and lateral diffusion of βArr2

To investigate the behavior of individual βArr2 and β_2_AR molecules on the plasma membrane of living cells, the two proteins were labeled with two distinct organic fluorophores via insertion of Halo- (Los et al., 2008) and SNAP- (Keppler et al., 2003) tags, respectively (**Figure 1A**). SNAP-β_2_AR and βArr2-Halo constructs – both fully functional (Calebiro et al., 2013) (**Figure S1A, B**) – were transiently expressed at low physiological levels (plasma membrane densities of 0.47 ± 0.13 and 0.40 ± 0.18 molecules/μm^2^, respectively) in Chinese Hamster Ovary (CHO) cells, which have no detectable β_1_/β_2_-ARs (Calebiro et al., 2013). The cells were then labeled with saturating concentrations of both dyes (**Figure S1C**) and simultaneously imaged by fast multicolor total internal reflection fluorescence (TIRF) microscopy combined with single-particle tracking (Sungkaworn et al., 2017) (**Figure 1B** and **Videos S1 and S2**); unspecific labeling was <1% (**Figure S1D**). We additionally visualized CCPs by co-transfection of GFP-labeled clathrin light chain. Data were acquired both under basal conditions and after early (2-7 min) and late (8-15 min) stimulation with the full β-adrenergic agonist isoproterenol. An excess of 2.4 million individual molecular trajectories were acquired and analyzed (**Table S1**). The results revealed that βArr2 molecules stochastically translocate from the cytoplasm to the plasma membrane, resulting in their sudden appearance in the TIRF field (**Figure 1C**). Unexpectedly, the newly translocated βArr2 molecules were often observed to diffuse on the plasma membrane for several frames before disappearing. Since photobleaching occurred on a much longer time scale (τ ~52 s) (**Figure S1E**), their disappearance was mostly due to dissociation from the plasma membrane. Of note, the majority (> 95%) of βArr2 molecules appeared at sites on the plasma membrane that were not occupied by β_2_ARs (**Figure 1D**). Although we observed accumulation of βArr2 molecules on the plasma membrane after isoproterenol stimulation (**Figure S1B**) – recapitulating their behavior in ensemble measurements (**Figure S1A**), the rate of appearance of new βArr2 molecules on the plasma membrane was remarkably similar between basal and stimulated conditions (**Figure 1E**). In contrast, the agonist-induced accumulation of βArr2 molecules on the plasma membrane was largely due to an increase in their membrane-bound lifetime. Under basal conditions, this had at least one larger, fast component (τ_fast_ = 0.58 s) and one smaller, slow component (τ_fslow_ = 4.53 s) (**Figure 1F**). Isoproterenol caused a ~4-5-fold increase in the magnitude of the second component corresponding to slowly dissociating βArr2 molecules, while having no relevant effects on the τ values of either component (τ_fast_ = 0.65/0.64 s and τ_fslow_ = 4.26/4.21 s for early/late stimulation) (**Figure 1F**).

**Figure 1.**
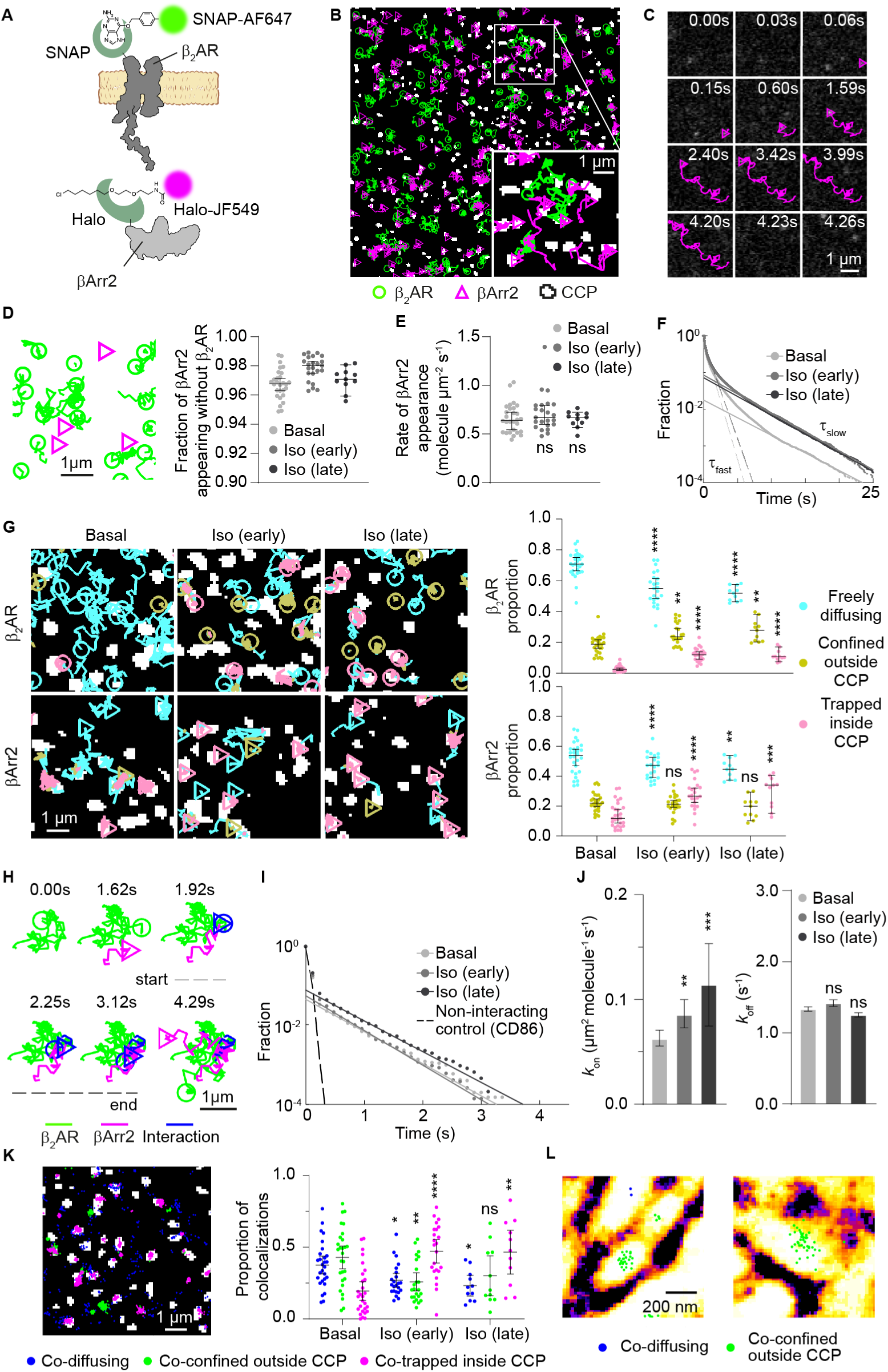
Single-molecule Imaging of I32AR and j3Arr2. (A) Labeling strategy. J3,AR was labeled with SNAP-AF647 via a SNAP-tag fused to its N-terminus. βArr2 was labeled with Halo-JF549 via a Halo-tag fused to its C-terminus. (B) Representative single-molecule results with trajectories overlaid on CCP binary mask. Inset, expanded view of the white box. (C) Representative trajectory of a βArr2 molecule appearing and transiently diffusing on the plasma membrane. (D) Sites of βArr2 appearance on the plasma membrane. Left, representative single-molecule trajectories showing the appearance of βArr2 molecules at sites unoccupied by receptors. Right, quantification. (E) Rates of βArr2 appearance on the plasma membrane. (F) Lifetimes of βArr2 molecules at the plasma membrane. Shown are the two main components estimated by exponential fitting. (G) Diffusivity states of β_2_AR (top) and βArr2 (bottom) molecules. Left, representative trajectories in cells with/without isoproterenol (10 μM) stimulation over CCP binary masks. Trajectory segments were assigned to three diffusivity states based on the results of a spatial confinement analysis and their colocalization with CCPs. Right, corresponding distributions. (H) Transient (~1.2 s) single-molecule colocalization event between a β_2_AR and a βArr2 molecule diffusing on the plasma membrane. (I) Relaxation curves of β_2_AR–βArr2 interactions, based on deconvolution of apparent colocalization times. CD86 was used as a non-interacting control. (J) Estimated *k*_on_ and *k*_off_ of β_2_AR–βArr2 interactions. (K) Representative spatial map (left) and overall distributions (right) of β_2_AR–βArr2 colocalization events in cells stimulated with isoproterenol (10 μM; late), color-coded based on the diffusivity states of the involved molecules. (L) β_2_AR–βArr2 single-molecule colocalizations over super-resolved (SRRF) image of actin filaments. Iso, isoproterenol. Early, 2-7 min. Late, 8-15 min. Data are median ± 95% confidence interval. n = 28, 23 and 11 cells for basal, Iso (early) and Iso (late), respectively. Differences in G, J (*k*_on_) and K are statistically significant by Kruskal Wallis test. *P < 0.05, **P < 0.01, ***P < 0.001, ****P < 0.0001 versus basal by t-test with Bonferroni correction. ns, statistically not significant. Images (B, C, G, K, L) are representative of a minimum of 3 independent experiments.

To gain insights into the diffusion properties of β_2_AR and βArr2 molecules on the plasma membrane, the obtained single-molecule trajectories were initially evaluated by time-averaged mean squared displacement (TAMSD) analysis (**Figure S2**). Since both β_2_AR and βArr2 molecules appeared to alternate phases of confinement and diffusion, we then used a new approach (Lanoiselée et al., 2021) to analyze these phases separately (**Figure S3A, B**). Under basal conditions, βArr2 molecules spent 52.3 ± 10.7% of their time freely diffusing, 22.6 ± 6.2% confined outside CCPs and 14.0 ± 8.3% trapped in CCPs (**Figure 1G**). By comparison, the corresponding values for β_2_AR were 70.5 ± 8.2%, 18.5 ± 6.0% and 2.9 ± 1.9% (**Figure 1G**). Isoproterenol stimulation caused ~4-fold and ~2-fold increases in the frequency of β_2_AR and βArr2 molecules trapped in CCPs and a modest increase (~1.4-fold) in that of β_2_ARs confined outside CCPs (**Figure 1G**). The durations of individual diffusion states were similar among all conditions, except for βArr2 trapping inside CCPs, which increased after agonist stimulation (**Figure S3C**).

Overall, these results unexpectedly revealed that βArr2 is in equilibrium between a cytosolic and a membrane-bound fraction, with βArr2 molecules spontaneously translocating to the plasma membrane under basal conditions and exploring space on the lipid bilayer via lateral diffusion. Moreover, they indicated that agonist stimulation prolongs the lifetime of βArr2 at the plasma membrane.

### β_2_AR–βArr2 interactions at the plasma membrane are highly dynamic and occur via lateral diffusion

We then asked how β_2_ARs–βArr2 interactions at the plasma membrane occur and for how long they last, which is presently unknown, but generally assumed to be sufficient for receptor– β-arrestin complexes to reach CCPs without dissociating (Laporte et al., 1999; Oakley et al., 1999).

Surprisingly, we found that most single-molecule β_2_ARs–βArr2 colocalizations were highly transient and often involved laterally diffusing β_2_AR and βArr2 molecules (**Figure 1H**). To estimate the frequency and duration of the underlying interactions, we applied our previously developed method based on deconvolution of apparent colocalization times with those of random colocalizations (Sungkaworn et al., 2017) (**Figure 1I**). Random colocalization times were estimated by imaging βArr2 with the unrelated integral membrane protein CD86 (Sungkaworn et al., 2017), which does not interact with GPCRs or βArr2 (expressed at levels comparable to β_2_AR, i.e. 0.53 ± 0.11 molecules/μm^2^). We estimated that β_2_AR–βArr2 interactions occurred with an association rate constant (*k*_on_) of 0.062 μm^2^ molecule^−1^ s^−1^ (95% confidence interval: 0.056–0.071) and lasted on average only ~0.7 s (dissociation rate/*k*_off_ = 1.34 s^-1^; 95% confidence interval: 1.31–1.37) in the absence of agonist (**Figure 1J**). Isoproterenol stimulation caused ~1.4 and ~1.8-fold increases in *k*_on_ at early and late time points, respectively, whilst causing no significant changes in *k*_off_ (**Figure 1J**).

We also investigated the location of β_2_AR–βArr2 interactions on the plasma membrane and whether they involved confined or freely diffusing molecules. Under basal conditions, β_2_AR–βArr2 colocalizations mainly involved freely diffusing molecules (37%) or molecules confined outside CCPs (47%) (**Figure 1K**). The latter could be at least partially explained by co-trapping in small nanodomains defined by the actin cytoskeleton (**Figure 1L**) as in the case of receptor– G protein interactions (Sungkaworn et al., 2017). As expected, isoproterenol stimulation increased (~3-fold) the proportion of single-molecule colocalization events between trapped molecules in CCPs (**Figure 1K**).

These results revealed that β_2_AR–βArr2 interactions mainly occur via lateral diffusion and are highly transient, lasting much shorter than generally assumed. Of note, the measured interaction times were considerably shorter than the average lifetime on the plasma membrane of the slowly dissociating component of βArr2 molecules that increased after agonist stimulation (**Figure 1F**). Moreover, they indicated that agonist stimulation controls β_2_ARs– βArr2 interactions primarily by increasing the rate of formation of productive receptor–β-arrestin complexes, rather than increasing their duration, similar to what we previously observed for receptor–G protein interactions (Calebiro et al., 2020; Sungkaworn et al., 2017).

### Receptors with varying affinity for βArr2 show different association rates but similar dissociation rates

Classical studies, comparing different GPCRs and chimeric receptors, identified two main classes of GPCRs based on their interactions with β-arrestins and trafficking properties (Oakley et al., 1999; Oakley et al., 2000). Class A GPCRs, such as the β_2_ARs, bind β-arrestins relatively weakly and appear to dissociate from β-arrestins during internalization. In contrast, Class B GPCRs have been suggested to bind β-arrestins more strongly and co-internalize with receptors in endosomes where they are seen colocalizing for extended periods of time (Oakley et al., 2000). Typical examples of Class B GPCRs are the vasopressin V2 receptor or the β_2_V_2_ chimeric receptor (Oakley et al., 1999; Shukla et al., 2014), which carries the C-tail of the vasopressin V2 receptor fused to the β_2_AR core.

To explore possible differences among receptors, we compared the β_2_AR with two additional GPCRs, the β_1_AR, which has an even weaker interaction with βArr2 than β_2_AR, and the β_2_V_2_ receptor, widely used as a model of strong βArr2 association (Shukla et al., 2014; Thomsen et al., 2016) (**Figure 2A**). All receptors showed similar lateral diffusion on the plasma membrane (**Figure S4A–C**). Agonist-dependent increases in βArr2 binding to both the plasma membrane and receptors, monitored by real-time bioluminescence resonance energy transfer (BRET) measurements, followed the expected order of β_2_V_2_>β_2_AR>β_1_AR (**Figure 2B**). Single-molecule imaging revealed a stronger accumulation of β_2_V_2_ than β_2_AR in CCPs after isoproterenol, with only a minor increase for β_1_AR (**Figure 2C** and **Figure S4B**). βArr2 accumulation in CCPs followed the same order of the receptors, albeit with higher basal and stimulated levels (**Figure 2C** and **Figure S4B**), indicating that receptor and β-arrestin accumulation are not stoichiometric.

**Figure 2.**
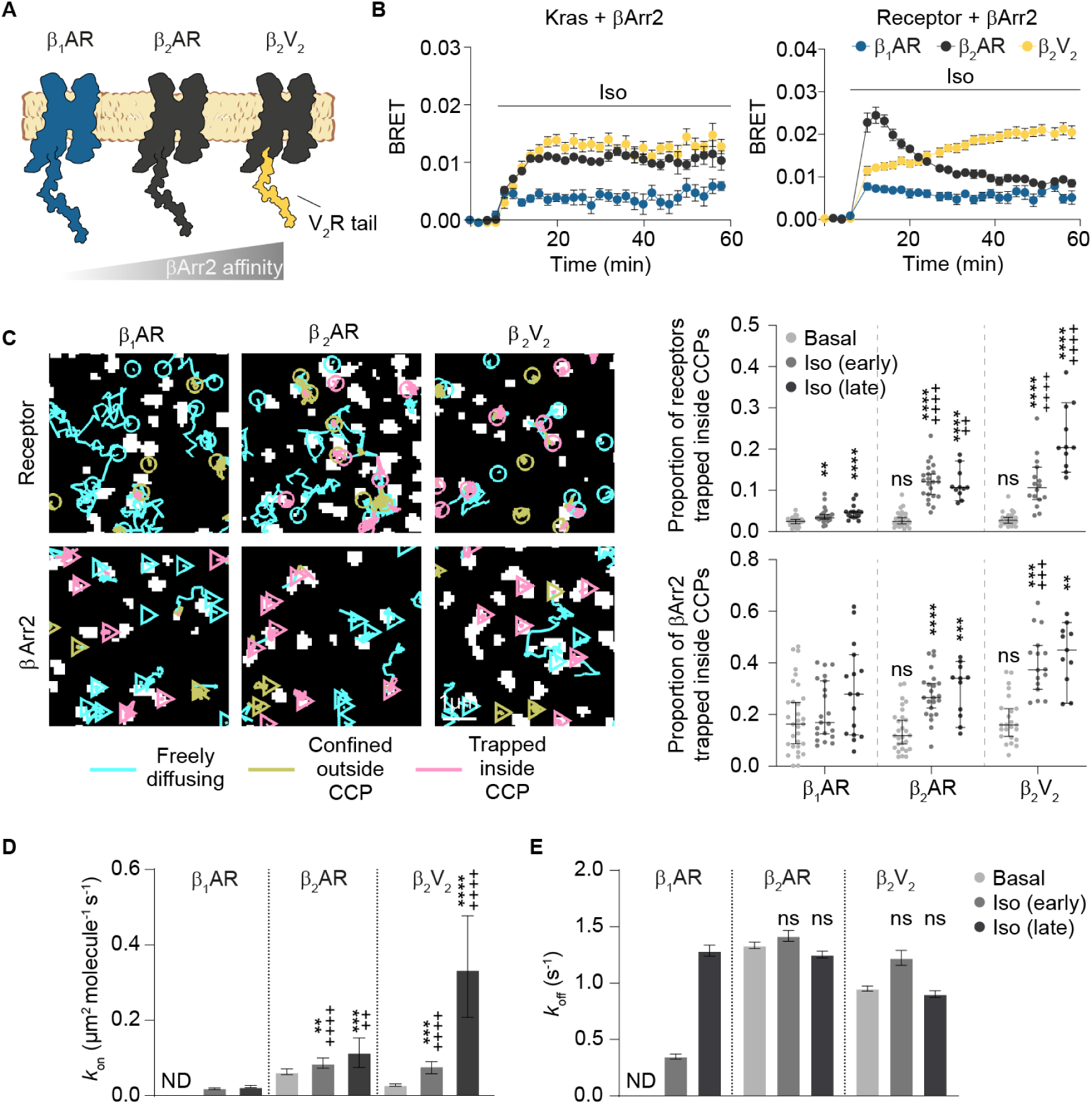
Affinity for Receptor C-Tail Governs βArr2 Interaction with Receptors and Plasma Membrane Behavior. (A) Schematic of the three investigated receptors (β_1_AR, β_2_AR and β_2_V_2_). (B) Kinetics of βArr2 recruitment to the plasma membrane and receptor upon isoproterenol (10 μM) stimulation. Shown are the results of real-time measurements monitoring BRET between NLuc fused to the plasma membrane marker Kras (left) or the receptor C-termini (right) and βArr2-Halo labeled with Halo-R110. (C) Diffusivity states of receptor (top) and βArr2 (bottom) molecules. Left, representative trajectories of individual receptor and βArr2 molecules in cells stimulated with isoproterenol (10 μM; late). Trajectory segments were assigned to three diffusivity states as in Figure 1G. Right, corresponding proportions of molecules trapped inside CCPs. (D, E) Estimated *k*_on_ (D) and *k*_off_ (E) values for β_1_AR, β_2_AR and β_2_V_2_–βArr2 interactions. Iso, isoproterenol. Early, 2-7 min. Late, 8-15 min. Data are mean ± s.e.m. (B), and median ± 95% confidence interval (C–E). n = 3 biological replicates (B), and 31, 21, 16, 28, 23, 11, 24, 16, 11 cells (C–E) for β_1_AR basal, β_1_AR Iso (early), β_1_AR Iso (late), β_2_AR basal, β_2_AR Iso (early), β_2_AR Iso (late), β_2_V_2_ basal, β_2_V_2_ Iso (early), β_2_V_2_ Iso (late), respectively. Results in C and D are statistically significant by Kruskal Wallis test. **P < 0.01, ***P < 0.001, ****P < 0.0001 versus corresponding basal condition, and ^++^P < 0.01, ^+++^P < 0.001, ^++++^P < 0.0001 versus corresponding β_1_AR condition by t-test with Bonferroni correction. ns, statistically not significant. Images in C are representative of a minimum of 3 independent experiments.

We then compared the kinetics of βArr2 interactions with the three receptors, estimated by deconvolution (**Figure 2D, E**). Basal β_1_AR–βArr2 interactions were undistinguishable from those with the CD86 control, consistent with the known low basal affinity of β_1_AR for βArr2. In contrast, we detected basal β_2_V_2_–βArr2 interactions (*k*_on_ 0.028 μm^2^ molecule^−1^ s^−1^; 95% confidence interval: 0.031–0.025) as in the case of β_2_AR. Isoproterenol increased *k*_on_ for all three receptors (β_1_AR ND; β_2_AR ~1.8-fold; β_2_V_2_ ~12-fold; **Figure 2D**), in good agreement with their affinity for βArr2. Relatively smaller differences were observed amongst *k*_off_ values (**Figure 2E**), with estimated average interaction times around ~1 s for all conditions, except for β_1_AR after early stimulation (~2.9 s).

As for β_2_AR, basal colocalization between β_1_AR or β_2_V_2_ and βArr2 molecules mainly involved free/confined molecules outside CCPs, with a variable increase inside CCPs after isoproterenol (**Figure S5**).

These results revealed that the interaction of three representative GPCRs with βArr2 is mainly controlled by their association rate (*k*_on_) and that receptor–β-arrestin interactions are short-lived, even in the case of the strongly interacting (Class B) β_2_V_2_ receptor.

### Sequence of events

To further dissect the sequence of events involved in β_2_AR–βArr2 interactions, we assigned to each molecule at each frame a state out of 6 unique ones (R1–6 and A1–6 for receptor and arrestin, respectively) defined by their motion (free/confined), mutual colocalization (present/absent) and trapping at CCPs (present/absent). A dummy state (R0/A0) was added to represent molecules before/after their appearance/disappearance from the plasma membrane (**Figure 3A and Table S2**). This information was used to build Markov chains describing both receptor and β-arrestin state occupancies and transitions (**Figure 3A**). Under basal conditions, β_2_ARs were prevalently exchanging between states corresponding to free diffusion (R1) and confinement outside CCPs (R3), with a small fraction trapped in CCPs alone (R5) (**Figure 3A**). Only minor fractions were co-diffusing with βArr2 molecules (R2), co-confined outside CCPs (R4) or co-trapped in CCPs (R6). βArr2 showed a similar pattern, albeit with a relatively higher proportion of molecules trapped in CCPs (A5) (**Figure 3A**). Isoproterenol stimulation increased the proportion of molecules trapped alone in CCPs by 3.6- and 1.7-fold, for β_2_AR and βArr2, respectively (**Figure 3B**). Unexpectedly, however, the transition from co-diffusion (R2, A2) to co-trapping in CCPs (R6, A6) was remarkably low under all conditions (**Figure 3A**). Instead, the main transition leading to co-trapping in CCPs was from the states corresponding to either molecule trapped alone in CCPs (R5, A5) (**Figure 3A**). We then focused on the sequence of states preceding and following single-molecule β_2_AR– βArr2 colocalizations (**Figure S6A**). We found that the majority of βArr2 molecules (86% at early stimulation) were laterally diffusing on the plasma membrane (A1) prior to co-diffusing (A2). Only ~16% of the colocalization events involved βArr2 molecules appearing from the cytoplasm (A0). Most (89%) βArr2 molecules continued to diffuse on the plasma membrane (A1) after co-diffusing with a receptor. Contrary to expectations, the state corresponding to βArr2 co-trapped with β_2_AR in CCPs (A6) was often preceded by βArr2 diffusing on the plasma membrane alone (A1), and we only rarely observed β_2_AR and βArr2 molecules reaching CCPs together (A2) (~2%) (**Figure S6B**). Remarkably, we also directly observed βArr2 molecules that continued diffusing on the plasma membrane after a transient β_2_AR–βArr2 colocalization until they reached and became trapped in a CCP alone (**Figure 3C**). Moreover, we observed βArr2 molecules visiting multiple CCPs via lateral diffusion (**Figure 3D**), indicating that βArr2 trapping at CCPs is reversible.

**Figure 3.**
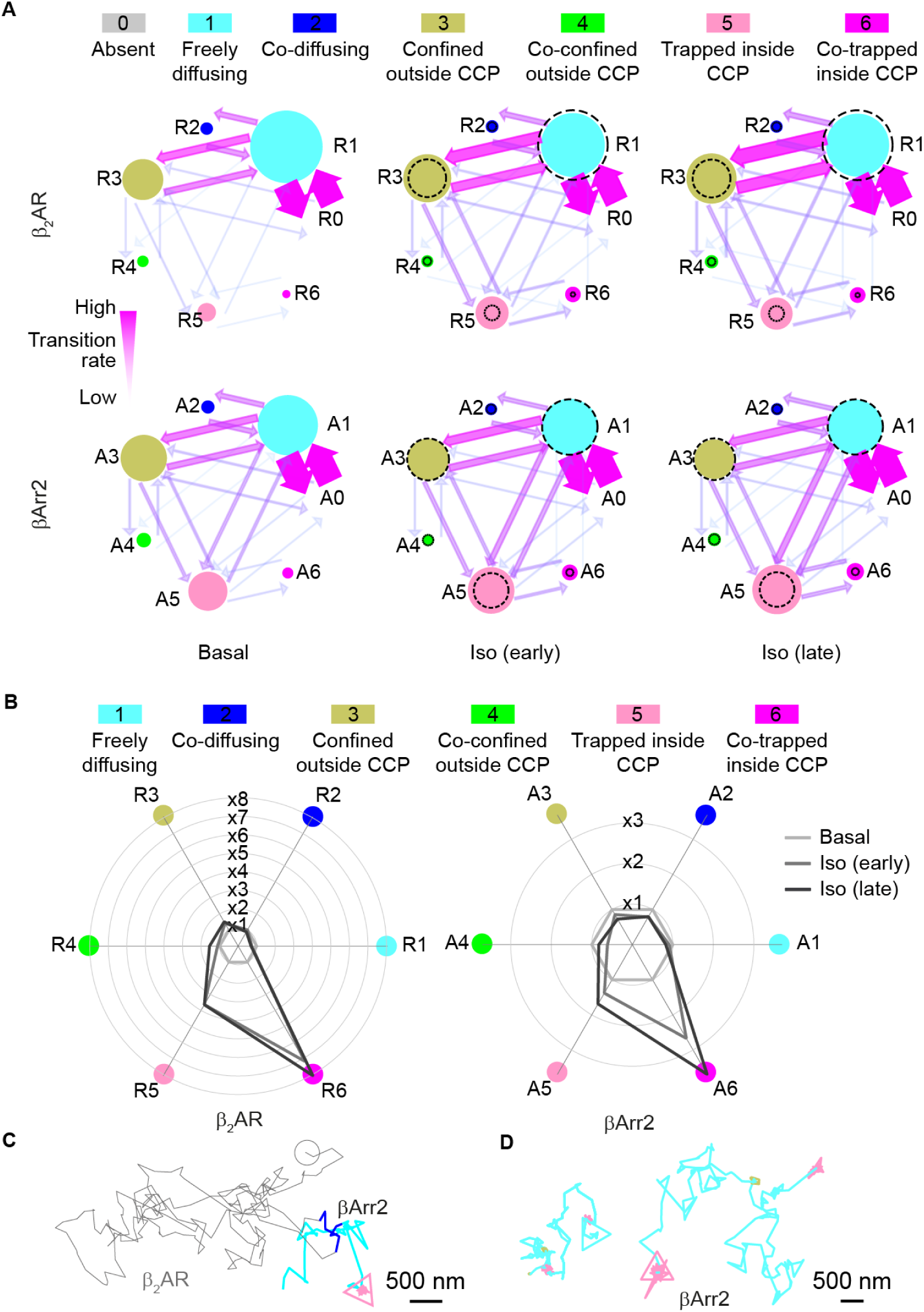
Sequence of Events in β_2_AR–βArr2 Interactions. For each frame, β_2_AR and βArr2 molecules were assigned to a unique state based on their mutual interactions and diffusion properties. (A) Markov chains showing relative state occupancies and forward transition probabilities for β_2_AR and βArr2 under basal, Iso (early) and Iso (late) conditions. Dashed circles, corresponding basal occupancies. (B) Radar plots showing the changes in state occupancies induced by isoproterenol (10 μM) stimulation. Data are normalized to the corresponding basal levels. (C) Example of a βArr2 molecule undergoing a transient interaction with a β_2_AR molecule to then reach a CCP without an accompanying receptor. The βArr2 trajectory is color-coded according to the identified states. (D) Examples of βArr2 molecules visiting multiple CCPs via lateral diffusion. Trajectories are shown as in C. Iso, isoproterenol. Early, 2-7 min. Late, 8-15 min. n = 88,851/51,147, 52,658/58,254, 27,741/22,769 transitions for β_2_AR/βArr2 basal, β_2_AR/βArr2 Iso (early), β_2_AR/βArr2 Iso (late), respectively.

Similar results, albeit with some quantitative differences, were observed for β_1_AR and β_2_V_2_ (**Figure S7A, B**). Remarkably, even in the case of the strongly interacting β_2_V_2_ receptor after isoproterenol stimulation, the vast majority of βArr2 molecules (83 %) reached CCPs alone. These results unexpectedly indicated that, after transient interaction, receptor and β-arrestin molecules mostly reach CCPs separately via lateral diffusion and not via co-diffusion as long-lived complexes.

### βArr2 spontaneously inserts into the lipid bilayer

Based on recent structural data on receptor–β-arrestin complexes in lipid nanodiscs (Lee et al., 2020; Staus et al., 2020) and biophysical measurements on visual arrestin (Lally et al., 2017), we hypothesized that the β-arrestin molecules seen spontaneously translocating to the plasma membrane and laterally diffusing without a receptor might be directly bound to the lipid bilayer via hydrophobic interactions. To test this hypothesis, we performed MD simulations starting with βArr2 close to the plasma membrane. As expected, βArr2 had a high degree of freedom in the cytoplasm assuming a wide range of positions relative to the plasma membrane in our simulations (**Figure 4A**). Remarkably, in 4 out of 40 independent simulations we observed spontaneous insertion of the βArr2 C-edge into the lipid bilayer (**Figure 4B** and **Table S3**), which restricted βArr2 in an overall orientation similar to the one found in receptor– β-arrestin complexes. In additional simulations starting from βArr2 with the C-edge inserted into the lipid bilayer, this was surprisingly followed by membrane penetration of the finger loop (**Figure 4B**), a key β-arrestin region involved in interaction with the receptor core (Cahill et al., 2017; Shukla et al., 2014). Based on extended MD simulations, we further refined a major lipid anchoring site in the C-edge (Leu192, Met193, Asp195, Arg332, Gly334), as well as two additional sites in the finger loop (Val71, Leu72, Gly73) and C-loop (Phe245, Ser246, Thr247, Ala248) (**Figure 4C**).

**Figure 4.**
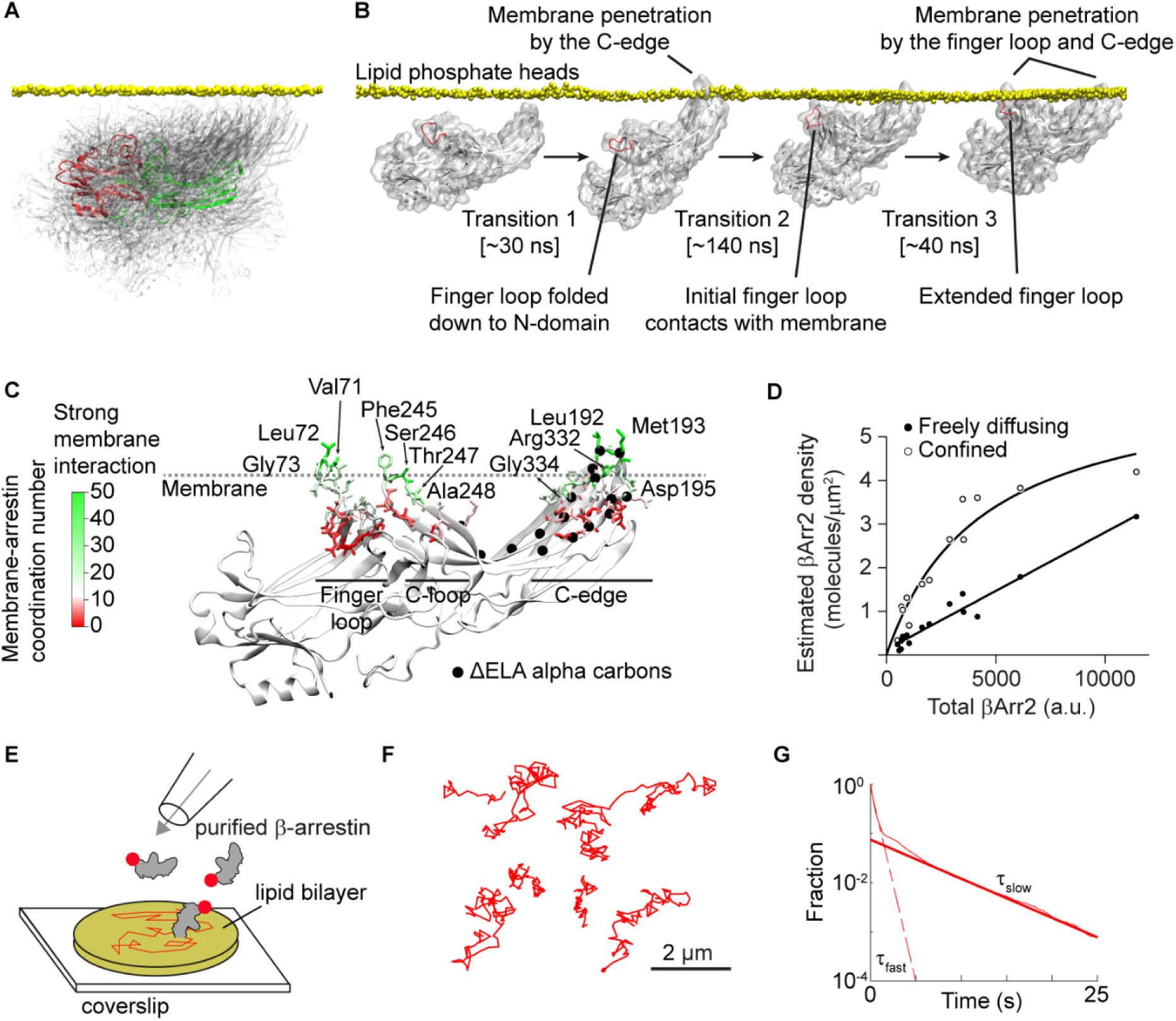
Spontaneous βArr2 Insertion into the Lipid Bilayer. (A) Superposition of the βArr2 conformations with different orientations relative to the plasma membrane (grey) sampled during an accumulated simulation time of 2.4 μs (time interval = 40 ns). A selected conformation is highlighted to show the position of the N- (red) and C- (green) domains. (B) Molecular dynamics (MD) simulation showing spontaneous insertion of the βArr2 C-edge into the lipid bilayer followed by a conformational rearrangement of the finger loop and its penetration into the bilayer. (C) Extended MD simulations (3 µs accumulated time) of membrane-bound βArr2. The results are shown on a representative structure of βArr2 after membrane insertion obtained in the simulations. Color indicates the lipid coordination numbers of βArr2 residues (see Methods). Main interacting residues (defined as those with a lipid coordination number > 20) in the C-edge as well as potential ones in the finger loop and C-loop are labeled. The amino acids mutated in the ΔELA mutant are indicated by black spheres. (D) Densities of freely diffusing βArr2 molecules on the plasma membrane of cells in which βArr2 expression was varied ~ 25-fold. n = 15 cells. (E) Schematic of the *in vitro* reconstitution experiments with purified β-arrestin and supported lipid bilayers. (F) Representative single-molecule trajectories showing spontaneous insertion and lateral diffusion of purified βArr2 in supported lipid bilayers. (G) Lifetimes of purified βArr2 molecules on supported lipid bilayers. Data are displayed and analyzed as in Figure 1F. n = 318,356 trajectories from 27 image sequences.

In support of our hypothesis, single-molecule experiments in which βArr2 expression was varied ~25-fold showed no saturation of freely diffusing βArr2 molecules on the plasma membrane at high βArr2 expression, consistent with their binding to high abundance sites such as membrane lipids rather than low-abundance receptors. In contrast, saturation was observed for confined βArr2 molecules, used as a control (**Figure 4D**).

To provide further evidence for direct binding of β-arrestin to the lipid bilayer, we resorted to a reconstituted system consisting of fluorescently labeled, purified β-arrestin and supported lipid bilayers obtained from giant unilamellar vesicles (GUVs) (**Figure 4E**), which were imaged by single-molecule microscopy. Importantly, also in this reconstituted system, we observed spontaneous insertion and lateral diffusion of β-arrestin in the supported lipid bilayers (**Figure 4F**). Moreover, the lifetime of β-arrestin molecules on the supported lipid bilayers (τ_fast_ = 0.55 s; τ_slow_ = 5.56 s) (**Figure 4G**) was remarkably similar to that in living cells (**Figure 1F**). Altogether, these findings provided strong evidence that β-arrestin can spontaneously insert into the lipid bilayer and remain associated with it for periods of times sufficient to explore the plasma membrane via lateral diffusion.

### βArr2 membrane pre-association drives receptor–β-arrestin interactions

To further test the mechanisms and functional consequences of β-arrestin pre-association with the plasma membrane, we generated a βArr2 mutant (ΔPIP2) carrying substitutions of key basic amino acids (K233Q/R237Q/K251Q) that form a known phosphatidylinositol 4,5-bisphosphate (PIP2) binding site (Gaidarov et al., 1999; Milano et al., 2002). A second extended lipid anchoring deficient mutant (ΔELA) (R189Q/F191E/L192S/M193G/T226S/K227E/T228S/K230Q/K231E/K233Q/R237E/K251Q/K 325Q/K327Q/V329S/V330D/R332E) (**Figure 4C**) was designed based on the available structures and our MD simulations to additionally interfere with C-edge lipid interactions. The introduction of multiple mutations was required due to the extensive contacts of βArr2 with the lipid bilayer. MD simulations of the βArr2 ΔELA-mutant indicate that the mutations do not disturb the secondary structure pattern observed in wild-type βArr2 (**Figure S8A, B**).

Mutating the PIP2 binding site alone (ΔPIP2) interfered with agonist-dependent increases in βArr2 binding to β_2_AR as well as to the plasma membrane and slowed down its accumulation at CCPs (**Figure 5A**), consistent with a role of PIP2 in facilitating receptor–arrestin interactions (Huang et al., 2020). However, it did not alter the basal frequency of βArr2 molecules exploring space via lateral diffusion (**Figure 5B, C**). In contrast, the ΔELA mutant was not only largely defective in agonist-dependent translocation and CCP accumulation (**Figure 5A, D**), but also in its ability to pre-associate with and diffuse laterally on the plasma membrane (**Figure 5B, C**). These results were further supported by metadynamics MD simulations with wild-type βArr2, which revealed a low energy well at 2.5-nm distance from the lipid bilayer, corresponding to βArr2 with the C-edge inserted in the bilayer, which was lost in the case of the ΔELA mutant (**Figure S8C**).

**Figure 5.**
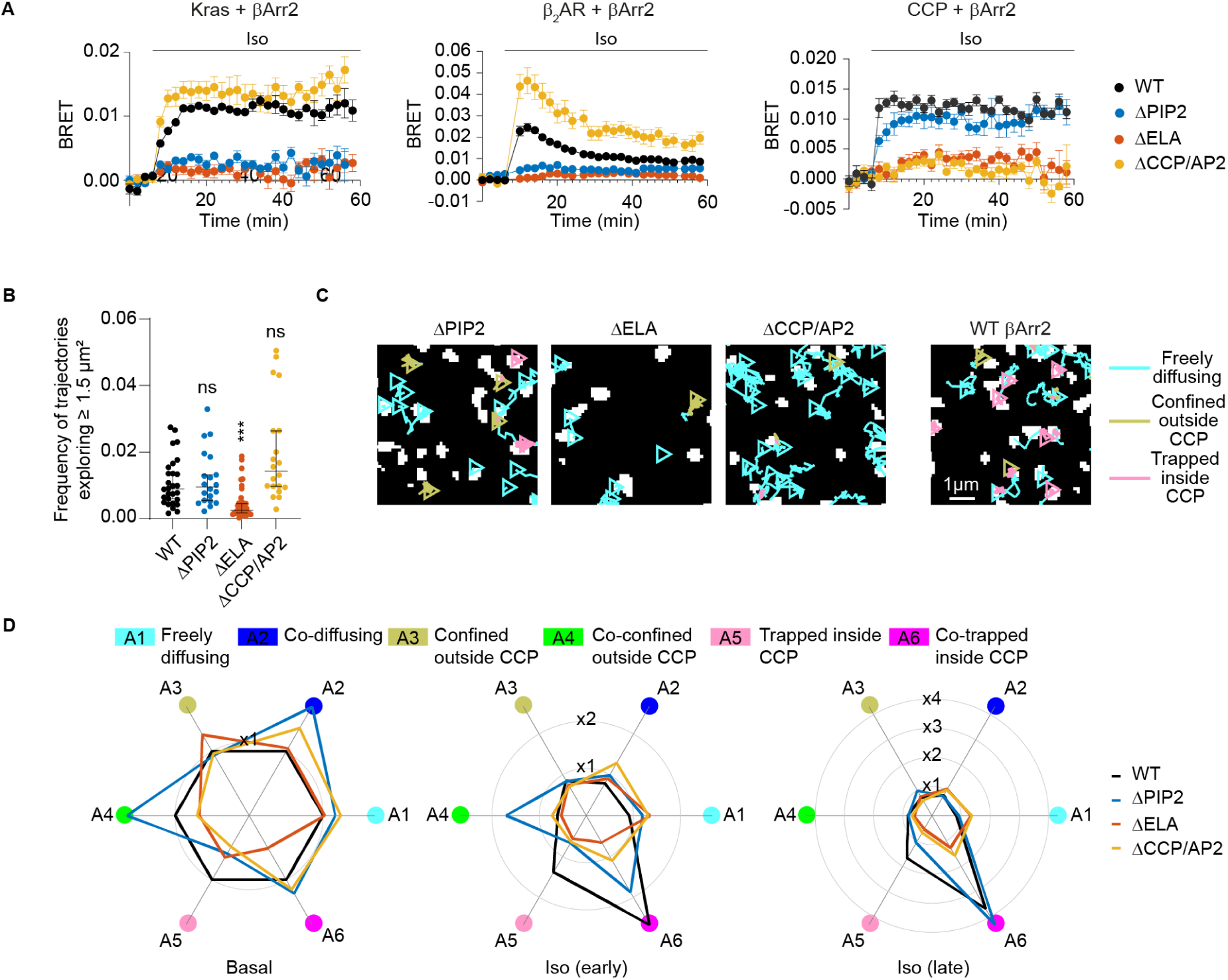
Functional Consequences of βArr2 Membrane Pre-association. (A) Kinetics of βArr2 mutant (ΔCCP/AP2, ΔPIP2 and ΔELA) recruitment to the plasma membrane (left), β_2_AR (middle), or CCPs (right) upon isoproterenol (10 μM) stimulation. Shown are the results of real-time measurements monitoring BRET between NLuc fused to the plasma membrane marker Kras (left), receptor C-terminus (middle), or clathrin light chain (right) and βArr2-Halo labeled with Halo-R110. (B) Propensity of βArr2 mutants to explore the plasma membrane. Shown are the relative frequencies of molecules exploring ≥ 1.5 μm^2^ in unstimulated cells. (C) Diffusivity states of βArr2 mutants. Shown are representative trajectories in cells stimulated with isoproterenol (10 μM; late). Results are shown as in Figure 1G. (D) Corresponding radar plots obtained from single-molecule experiments showing the changes in state occupancies induced by isoproterenol (10 μM) stimulation. Iso, isoproterenol. Early, 2-7 min. Late, 8-15 min. Wild-type (WT) βArr2 is included in A-D for comparison. Data are mean ± s.e.m. (A) and median ± 95% confidence interval (B). n = 3 biological replicates (A), 28, 20, 20, 34 cells (B) for WT basal, ΔCCP/AP2 basal, ΔPIP2 basal, ΔELA basal, and 51,147, 58,254, 22,769, 43,242, 37,370, 40,079, 38,095, 20,436, 30,530, 76,129, 18,683, and 27,349 transitions (D) for WT basal, WT Iso (early), WT Iso (late), ΔCCP/AP2 basal, ΔCCP/AP2 Iso (early), ΔCCP/AP2 Iso (late), ΔPIP2 basal, ΔPIP2 Iso (early), ΔPIP2 Iso (late), ΔELA basal, ΔELA Iso (early) and ΔELA Iso (late), respectively. Differences in B are statistically significant by Kruskal Wallis test. ***P < 0.001 versus wild type by t-test with Bonferroni correction. Images in C are representative of a minimum of 3 independent experiments.

Our hypothesis was further supported by experiments with a clathrin/AP2 binding-deficient βArr2 mutant (ΔCCP/AP2), which, instead of accumulating in CCPs, continued diffusing laterally on the plasma membrane after isoproterenol stimulation (**Figure 5A-D**). These results also ruled out the possibility that the laterally diffusing βArr2 molecules might be bound to the plasma membrane via clathrin/AP2.

Overall, these results support a model in which βArr2 binds directly to the plasma membrane with a major contribution of hydrophobic residues in the C-edge, whereas the previously identified PIP2 binding site appears dispensable for membrane anchoring.

### The β_2_AR C-tail mediates βArr2 activation

β-arrestin has been shown to interact with receptors via two distinct poses (**Figure 6A**), involving either interactions between a polar core in the β-arrestin N-domain and the receptor phosphorylated C-tail or between the β-arrestin finger loop and the receptor core (Cahill et al., 2017; Eichel et al., 2018; Shukla et al., 2014). To dissect the contribution of either modality, we studied β_2_AR constructs carrying a deletion in either the third intracellular loop (ΔICL3) or C-tail (ΔC-tail). Whereas the ΔC-tail truncation disrupts phosphorylation-mediated interaction with β-arrestin, the ΔICL3 deletion has been demonstrated to disrupt β-arrestin core-interaction using comprehensive bimane fluorescence spectroscopy analyses (Ghosh et al., 2019; Kumari et al., 2016; Kumari et al., 2017). The ΔICL3 mutant showed normal agonist-dependent increases in βArr2 interaction, βArr2 plasma membrane binding and accumulation together with βArr2 in CCPs (**Figure 6B-D**), consistent with previous ensemble measurements (Kumari et al., 2017). In contrast, the ΔC-tail mutant was unable to accumulate in CCPs and failed to promote agonist-dependent β_2_AR–βArr2 interactions, βArr2 translocation to the plasma membrane and βArr2 accumulation in CCPs (**Figure 6B-D**).

**Figure 6.**
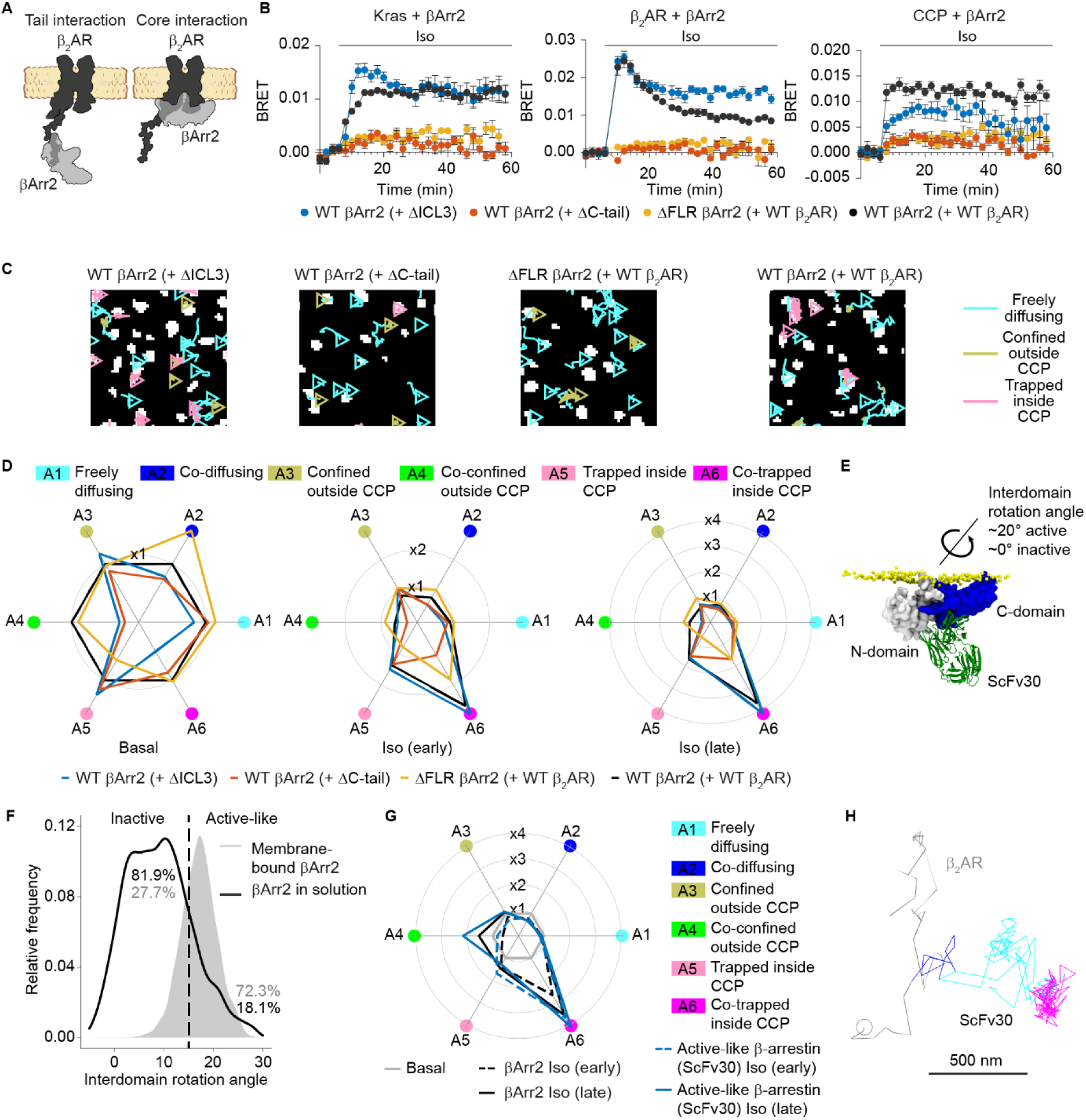
Mechanisms of βArr2 Activation and Stabilization at the Plasma Membrane. (A) Schematic of C-tail and core receptor–arrestin interactions. (B) Kinetics of βArr2 recruitment to the plasma membrane (left), β_2_AR (middle), or CCPs (right) in cells expressing the indicated combinations of wild-type (WT) or mutant βArr2 and β_2_AR constructs and stimulated with isoproterenol (10 μM). Results are shown as in Figure 5A. (C) βArr2 diffusivity states in cells expressing the same combinations of constructs. Shown are representative trajectories after stimulation with isoproterenol (10 μM; late). Cells expressing WT β_2_AR and βArr2 are included for comparison. Results are shown as in Figure 1G. (D) Corresponding radar plots obtained from single-molecule experiments showing the changes in state occupancies induced by isoproterenol (10 μM) stimulation. (E) Recognition of active-like membrane-bound βArr2 by Fab30/ScFv30. The structural model was obtained by aligning the membrane-bound βArr2 conformation obtained in MD simulations to the crystal structure of the βArr1–Fab30 complex (PDB 4JQI). (F) Results of MD simulations comparing βArr2 conformations in solution and bound to the lipid bilayer. The inter-domain rotation angle between the N- and C-domains is used as a measure of βArr2 activation. (G) Radar plot obtained from single-molecule experiments comparing the behavior of ScFv30, recognizing active-like β-arrestin, and total βArr2. (H) Example of an active-like β-arrestin molecule, visualized with ScFv30, undergoing transient interaction with a β_2_AR molecule (blue) to then diffuse away alone (cyan) and reach a CCP without the receptor (magenta). Iso, isoproterenol. Early, 2-7 min. Late, 8-15 min. Data are mean ± s.e.m. (B). n = 3 biological replicates (B), 51,147, 58,254, 22,769, 45,670, 41,068, 37,071, 50,911, 44,549, 38,586, 57,627, 64,695, 46,518 transitions (D) for WT basal, WT Iso (early), WT Iso (late), ΔICL3 basal, ΔICL3 Iso (early), ΔICL3 Iso (late), ΔC-tail basal, ΔC-tail Iso (early), ΔC-tail Iso (late), ΔFLR basal, ΔFLR Iso (early), ΔFLR Iso (late), and 64,757, 65,227, 99,428 transitions (G) for ScFv30 basal, ScFv30 Iso (early), ScFv30 Iso (late), respectively. Images in C are representative of a minimum of 3 independent experiments.

We then introduced a βArr2 mutant carrying a deletion in the finger loop (ΔFLR), which has also been shown to interfere with β-arrestin core interaction (Cahill et al., 2017). Although the ΔFLR was capable of binding to the plasma membrane and exploring space via lateral diffusion under basal conditions (frequency of molecules exploring ≥ 1.5 μm^2^ ~0.09, comparable to wild type), it was largely deficient in agonist-induced translocation to the plasma membrane and accumulation in CCPs (**Figure 6B-D**), in striking contrast to the ΔICL3 mutant. These results indicate that βArr2 engagement with the receptor C-tail but not its core is required for agonist-induced accumulation of both receptors and arrestins in CCPs. Moreover, they suggest that an intact finger loop is required for βArr2 activation and accumulation in CCPs.

### Lipid interactions stabilize active-like βArr2 on the plasma membrane

Our MD simulations revealed a previously unrecognized potential interaction of the finger loop with the plasma membrane (**Figure 4B, C**) and suggested that this interaction may promote and/or stabilize the finger loop in an active-like state. Since β-arrestin activation involves a key inter-domain rotation between its N- and C-domains, we used this as a measure of βArr2 activation (inactive-like < 15°, active-like > 15°) (Dwivedi-Agnihotri et al., 2020) (**Figure 6E**). Interestingly, we found that full βArr2 lipid anchoring (i.e., via finger loop, C-loop and C-edge) is associated with a shift of the inactive–active βArr2 equilibrium towards active-like conformations in MD simulations (**Figure 6F**).

To further test this hypothesis, we took advantage of an intrabody based on a single-chain variable fragment (ScFv30) that selectively recognizes the active-like rotation in βArr1/2 (Ghosh et al., 2019; Ghosh et al., 2017; Shukla et al., 2014) (**Figure 6E**). BRET experiments in βArr1/2 CRISPR/Cas9 knockout cells, in which we re-expressed βArr2, confirmed the ability of ScFv30 to recognize active-like βArr2, as shown by its plasma membrane translocation following isoproterenol stimulation, which did not occur in control knockout cells (**Figure S9A**). Remarkably, single-molecule imaging revealed striking similarities between the behavior of ScFv30 and βArr2, including the presence of laterally diffusing ScFv30 molecules on the plasma membrane with characteristics superimposable to those of βArr2 and capable of reaching and becoming trapped in CCPs without an accompanying receptor (**Figure 6G, H** and **Figure S9B-D**).

Altogether, these results strongly support a model in which transient interaction with an active receptor catalyzes the conversion of βArr2 into an active-like conformation that is stabilized on the plasma membrane via lipid interactions, prolonging βArr2 lifetime at the plasma membrane and allowing it to reach CCPs alone.

## DISCUSSION

According to the current model, which is largely based on ensemble measurements, β-arrestin is assumed to translocate from the cytoplasm to directly bind an active receptor on the plasma membrane and remain bound to the same receptor at least until they reach a CCP together. In contrast, our single-molecule results surprisingly reveal a completely different scenario whereby β-arrestin spontaneously pre-associates with the plasma membrane, *de facto* behaving like a membrane protein. This allows β-arrestin to explore space via lateral diffusion and undergo highly transient interactions with receptors that lead to β-arrestin activation. Importantly, this prolongs the lifetime of β-arrestin at the plasma membrane allowing it to reach CCPs independently of the initial, short-lived receptor–β-arrestin complex. These findings are surprising as, in several respects, they liken the behavior of β-arrestin to that of G proteins, the other main class of GPCR signaling partners.

Based on our detailed single-molecule measurements, we propose a new multistep model for receptor–β-arrestin interactions, which involves at least 5 distinct events: (i) β-arrestin spontaneously inserts into the plasma membrane via its C-edge, (ii) laterally diffuses on the plasma membrane, (iii) transiently (~ 1 s) interacts with an active receptor via lateral diffusion and becomes activated with extension of the finger loop, (iv) is stabilized in a membrane-bound, active-like conformation with involvement of the finger loop, and (v) diffuses to CCPs separately from the activating receptor (**Figure 7**).

**Figure 7.**
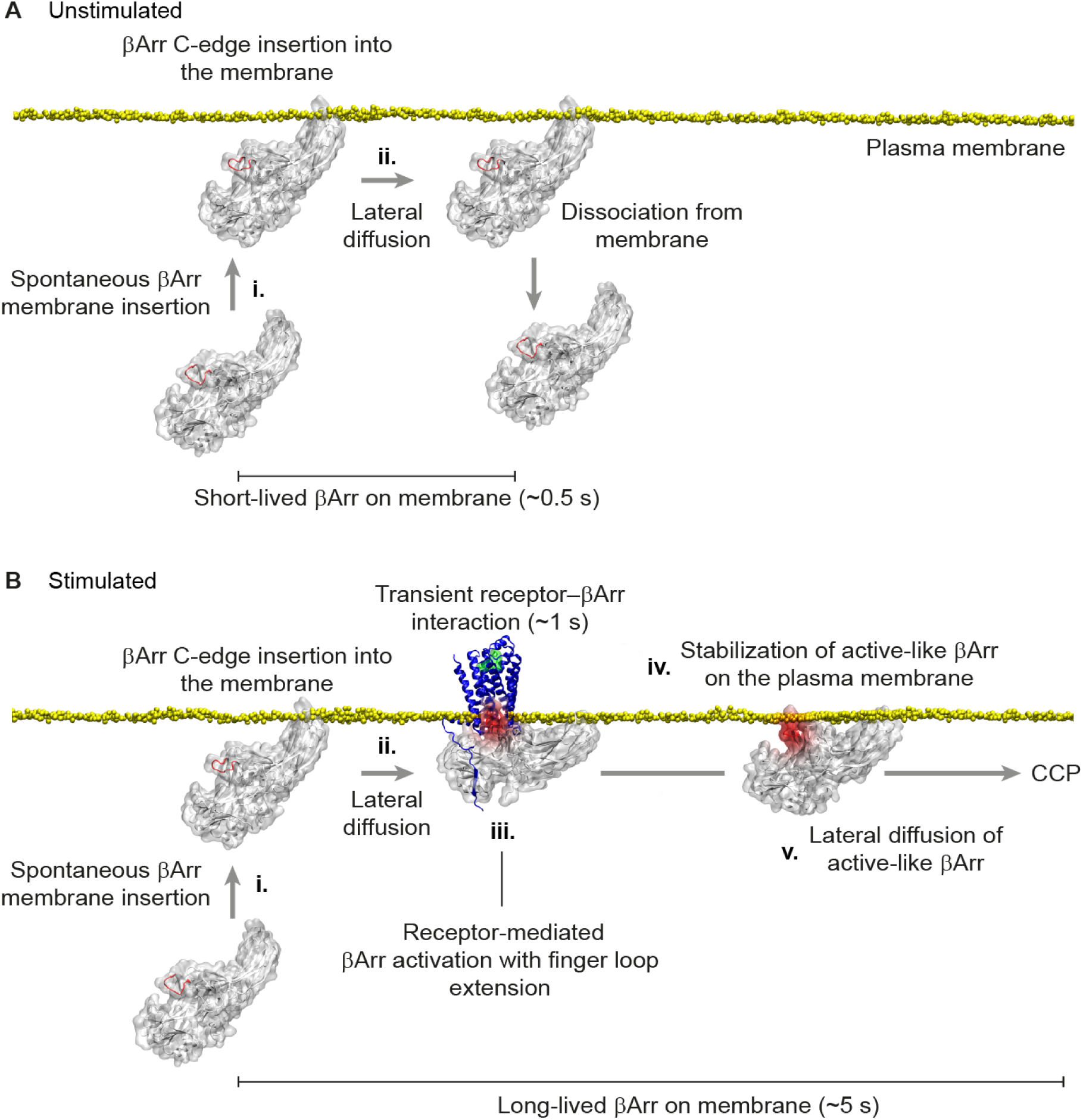
Proposed Model of Receptor–β-Arrestin Interactions. (A) Behavior of β-arrestin at the plasma membrane under unstimulated conditions. Inactive β-arrestin in the cytoplasm spontaneously binds to the plasma membrane via insertion of the C-edge into the lipid bilayer (i), allowing it to explore space via lateral diffusion (ii). Most β-arrestin molecules remain on the plasma membrane for a relatively short time before dissociating and returning to the cytoplasm. (B) Behavior of β-arrestin at the plasma membrane in the presence of a stimulated receptor. After spontaneous insertion into the plasma membrane (i), β-arrestin reaches the receptor via lateral diffusion (ii). Transient interaction with the phosphorylated receptor C-tail catalyzes β-arrestin activation, including β-arrestin interdomain rotation and extension of the finger loop (iii). Upon dissociation from the receptor, the interaction of the extended finger loop with the lipid bilayer contributes to stabilizing β-arrestin in a membrane-bound, active-like conformation (iv). This allows active-like β-arrestin to stay longer on the plasma membrane and reach CCPs vial lateral diffusion without an accompanying receptor (v).

A first key finding of our study is that β-arrestin spontaneously interacts with the lipid bilayer prior to meeting a receptor and that the latter happens via lateral diffusion. This reveals a previously unexpected role of the lipid bilayer in facilitating receptor–β-arrestin interactions, which likely occurs via at least three separate mechanisms. First, the pre-association of β-arrestin with the lipid bilayer via its C-edge restricts β-arrestin in an overall orientation relative to the plasma membrane that matches the orientation in receptor–β-arrestin complexes. Second, the pre-association of β-arrestin with the plasma membrane increases its local concentration close to the receptors. Third, the switch from 3D diffusion in the cytoplasm to 2D diffusion on the plasma membrane can reduce the time required to reach a receptor (Bénichou et al., 2010; Bénichou et al., 2011). All these factors likely concur to increase the speed and efficiency of receptor–β-arrestin interactions.

Another key finding of our study is that receptor–β-arrestin interactions on the plasma membrane are highly dynamic and short-lived. Whereas the duration of these key interactions in GPCR signaling was previously unknown, receptor–β-arrestin complexes were widely assumed to be sufficiently stable to allow receptors and β-arrestins to reach CCPs together, as in the case of Class A GPCRs, or even co-internalize and remain associated in endosomes, as in the case of Class B receptors (Oakley et al., 1999, 2001; Oakley et al., 2000). In contrast, our single-molecule results reveal that receptor–β-arrestin interactions are highly transient, lasting on average only approximately 1 s. This is not only true for β_1_AR and β_2_AR, but also for β_2_V_2,_ which has been previously reported to co-internalize and remain colocalized with β-arrestin in endosomes for at least one hour (Oakley et al., 1999). Our finding that receptor–β-arrestin interactions are highly transient has a number of important implications. First, it means that most receptors and β-arrestins must reach CCPs separately. Moreover, it implies that an average receptor meets multiple β-arrestin molecules during its lifetime on the plasma membrane, permitting a much more dynamic regulation of GPCR signaling than previously thought. This likely occurs at least two times. A first one when a receptor and a β-arrestin molecule transiently interact leading to β-arrestin activation, and a second one when receptors and β-arrestins meet again in CCPs.

Our results also show that receptor–β-arrestin interactions and the ensuing β-arrestin activation require an intact receptor C-tail, whereas interaction with the receptor core is dispensable, providing further support to previous results obtained with ensemble methods (Kumari et al., 2016; Kumari et al., 2017; Nobles et al., 2007; Shukla et al., 2013; Shukla et al., 2014; Xiao et al., 2004). This is distinct from a previously proposed mechanism for prolonged β-arrestin activation, which, instead, relied on interaction of β-arrestin with the receptor core for β-arrestin activation and ruled out the involvement of either the receptor C-tail or β-arrestin C-edge (Eichel et al., 2018; Latorraca et al., 2018). It is tempting to speculate that evolution might have produced two separate receptor interaction sites on β-arrestin to allow it to independently perform its two main functions, i.e., receptor desensitization, mediated by the finger loop–core interaction, and β-arrestin activation, mediated by the N-domain–C-tail interaction.

Moreover, our study identified the C-edge as the critical region for β-arrestin membrane anchoring as well as additional membrane interaction sites in the finger loop and C-loop. Our results suggest that finger loop extension and its subsequent lipid interaction play an important role in maintaining β-arrestin in an active-like conformation at the plasma membrane. Whereas this appears to occur spontaneously as suggested by our MD simulations and results with supported lipid bilayers, the interaction with an active receptor likely catalyzes this transition. This is consistent with the finding that interactions of proximal phosphate groups in the receptor C-tail can trigger the release of an ionic lock that keeps the finger loop in its inactive conformation (Sente et al., 2018). This, in turn, would allow the finger loop to interact with the plasma membrane stabilizing β-arrestin in a membrane-bound, active-like conformation, capable of reaching CCPs alone to mediate GPCR internalization and ‘non-classical’ signaling. This model is supported by our finding that agonist stimulation increases the lifetime of β-arrestin at the plasma membrane. Since we estimate that active-like β-arrestin dissociates from the plasma membrane much slower than inactive β-arrestin, this emerges as the main reason why β-arrestin accumulates on the plasma membrane after agonist stimulation as opposed to the formation of rather stable, long-lived receptor–β-arrestin complexes, as previously thought. Our findings also suggest an entirely new role for the finger loop in prolonging β-arrestin activity, which is distinct from its function in core-mediated signal desensitization.

At the same time, our results indicate that the known β-arrestin PIP2 binding site (Gaidarov et al., 1999; Milano et al., 2002), although dispensable for membrane anchoring, plays an important role in facilitating receptor–β-arrestin interactions, consistent with the recent finding of a PIP2 bridge in the structure of the neurotensin receptor 1–βArr1 complex (Huang et al., 2020).

Whereas previous studies investigated β-arrestin with various methods, the new insights obtained in this study were only possible thanks to our single-molecule approach, which allowed us to directly observe and dissect the precise sequence of events during the entire lifetime of individual β-arrestin molecules on the plasma membrane (from their translocation from the cytoplasm to accumulation in CCPs), which cannot be deduced from ensemble measurements.

Altogether, our findings redefine the current model of receptor–β-arrestin interactions by revealing a previously unrecognized, critical role of β-arrestin interactions with the lipid bilayer in mediating β-arrestin interactions with receptors via lateral diffusion and its accumulation on the plasma membrane in an active-like conformation.

## ACKNOWLEDGEMENTS

This study was supported by a Wellcome Trust Senior Research Fellowship (212313/Z/18/Z to D.C.). Research program in the laboratory of A.K.S. is supported by a Senior Fellowship of the DBT/Wellcome Trust India Alliance (IA/S/20/1/504916). A.K.S. is a EMBO Young Investigator and Joy Gill Chair Professor. We thank Chris Tate for discussions during the initial phase of this study and Ravi Mistry for technical support.

## AUTHOR CONTRIBUTIONS

D.C. conceived the study. J.G., Z.K., Y.L, T.M. and S.L.O.B. performed the single-molecule and BRET experiments and analyzed the data. Y.L. and D.C. developed the mathematical analyses and wrote the software. T.M.S. and B.M.L. performed the MD simulations. D.C. supervised the single-molecule experiments. S.H. and D.C. supervised the BRET experiments. J.S. supervised the MD simulations. M.B. and A.S. generated the ScFv30 probe and provided support with ScFv30 experiments. M.M. and D.M.O. prepared the GUVs and provided support with the experiments with supported lipid bilayers. D.C., J.G., Z.K. and Y.L. wrote the manuscript with contributions from J.S., T.M.S. and B.M.L. All authors discussed the results and edited the manuscript.

## SUPPLEMENTAL FIGURES

**Figure S1.**
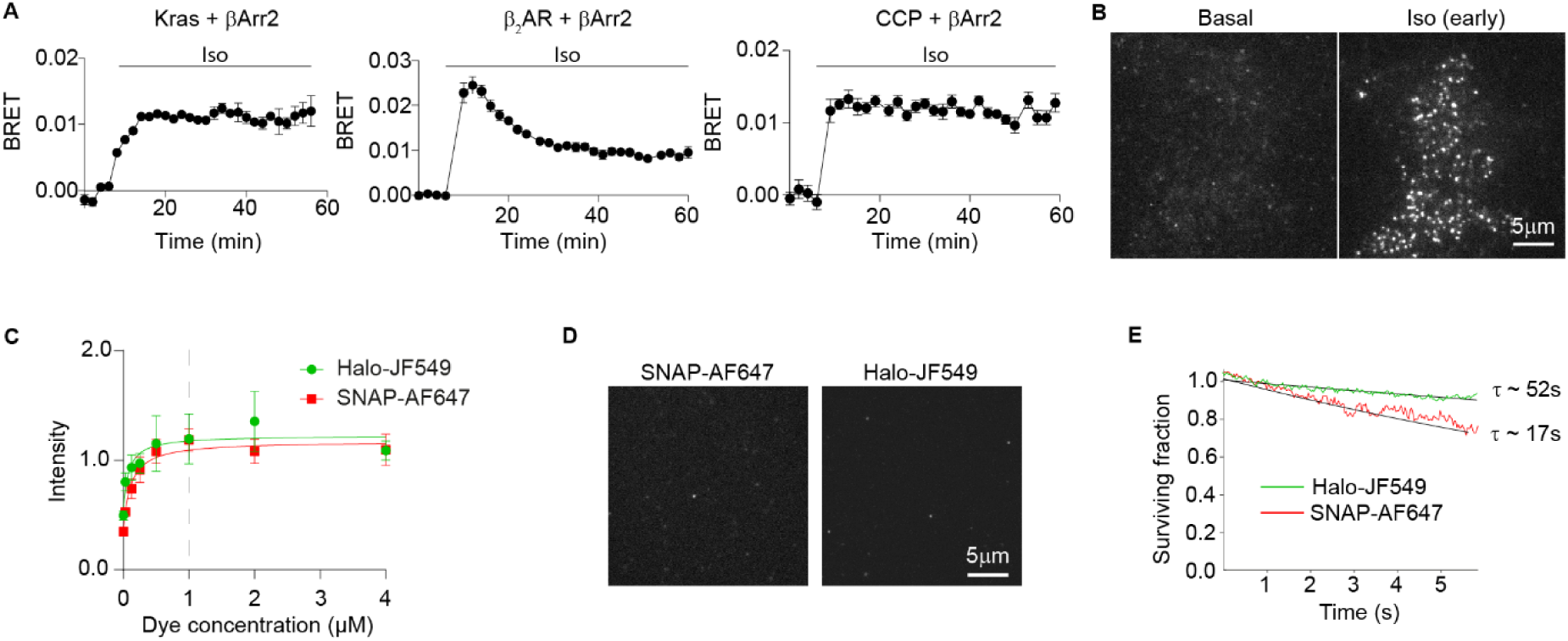
Control Experiments, Related to Figure 1. (A) Functional characterization of the Halo-tagged βArr2 construct. Shown are the results of real-time measurements monitoring BRET between NLuc fused to Kras (plasma membrane recruitment; left), β_2_AR (receptor binding; middle) or clathrin light chain (CCP recruitment; right) and βArr2-Halo labeled with Halo-R110. Cells were stimulated with isoproterenol (10 µM). (B) Representative single-molecule experiment showing rapid plasma membrane translocation of individual βArr2-Halo molecules upon stimulation with isoproterenol (10 µM). (C) Labeling efficiency. Shown are fluorescence intensity values in cells expressing β_2_AR carrying either an intracellular Halo tag or an extracellular SNAP-tag, incubated with increasing concentrations of Halo-JF549 or SNAP-AF647, respectively. Saturating concentrations of both dyes (1 μM) were subsequently used for single-molecule microscopy experiments. (D) Non-specific labeling. Mock-transfected cells were labeled with either SNAP-AF647 or Halo-JF549 and imaged by single-molecule microscopy. (E) Photobleaching rates. Shown are the photobleaching curves for Halo-JF549 and SNAP-AF647 obtained from single-molecule experiments. Iso, isoproterenol. Early, 2-7 min. τ, average lifetime. Data are mean ± s.e.m. (A) or mean ± s.d. (C). n = 4 replicates (A), 10 cells (C) and 28 cells (E). Images (B, D) are representative of 4 independent experiments.

**Figure S2.**
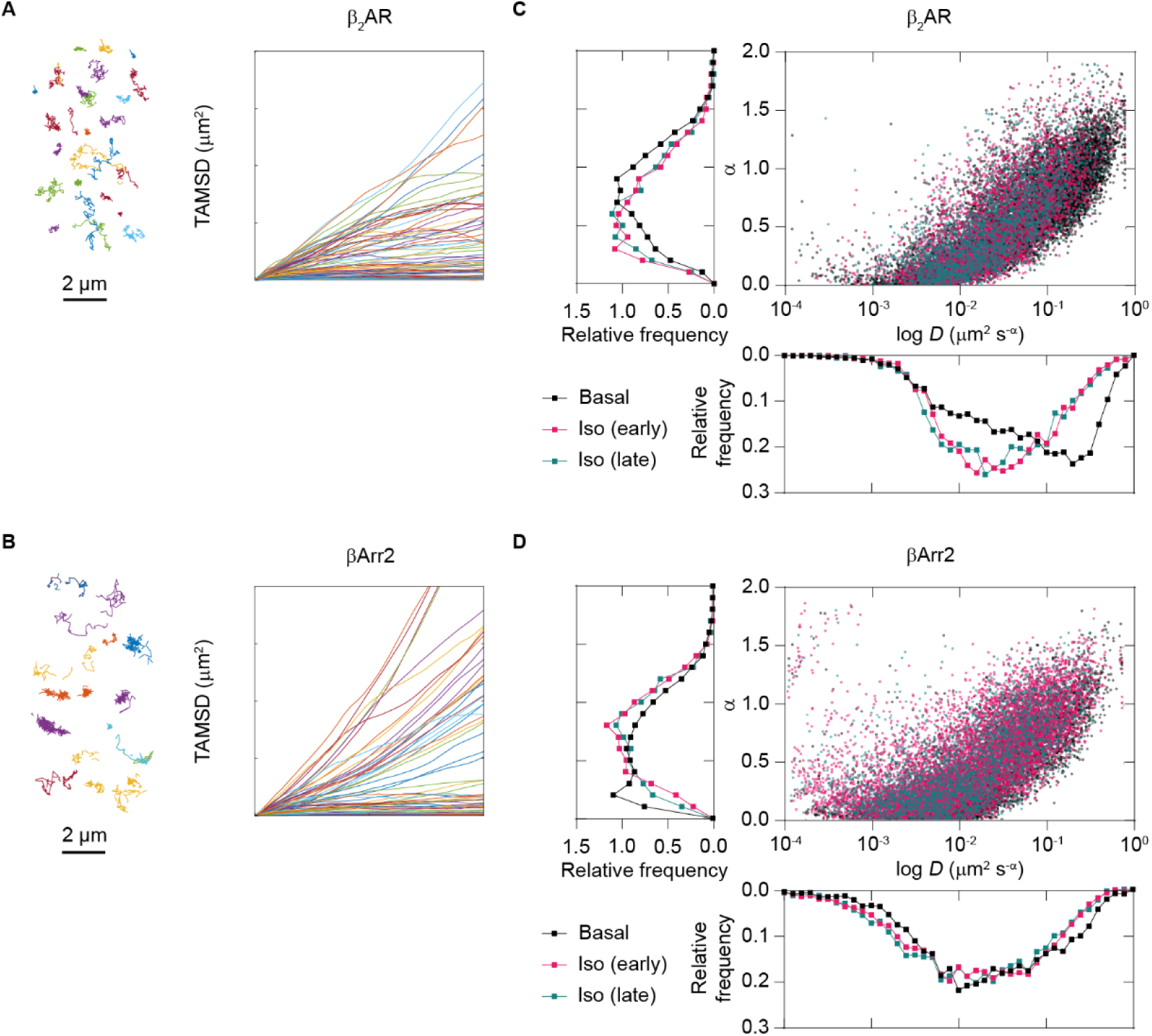
Time-Averaged Mean Squared Displacement (TAMSD) Analysis, Related to Figure 1. (A, B) Representative SNAP-β_2_AR (A) and βArr2-Halo (B) trajectories and corresponding TAMSD curves. (C, D) Scatter plots and distributions of diffusion coefficient (*D*) and anomalous diffusion exponent (α) values estimated for β_2_AR (C) and βArr2 (D) trajectories without (basal) or with isoproterenol (10 μM) stimulation. Iso, isoproterenol. Early, 2-7 min. Late, 8-15 min.

**Figure S3.**
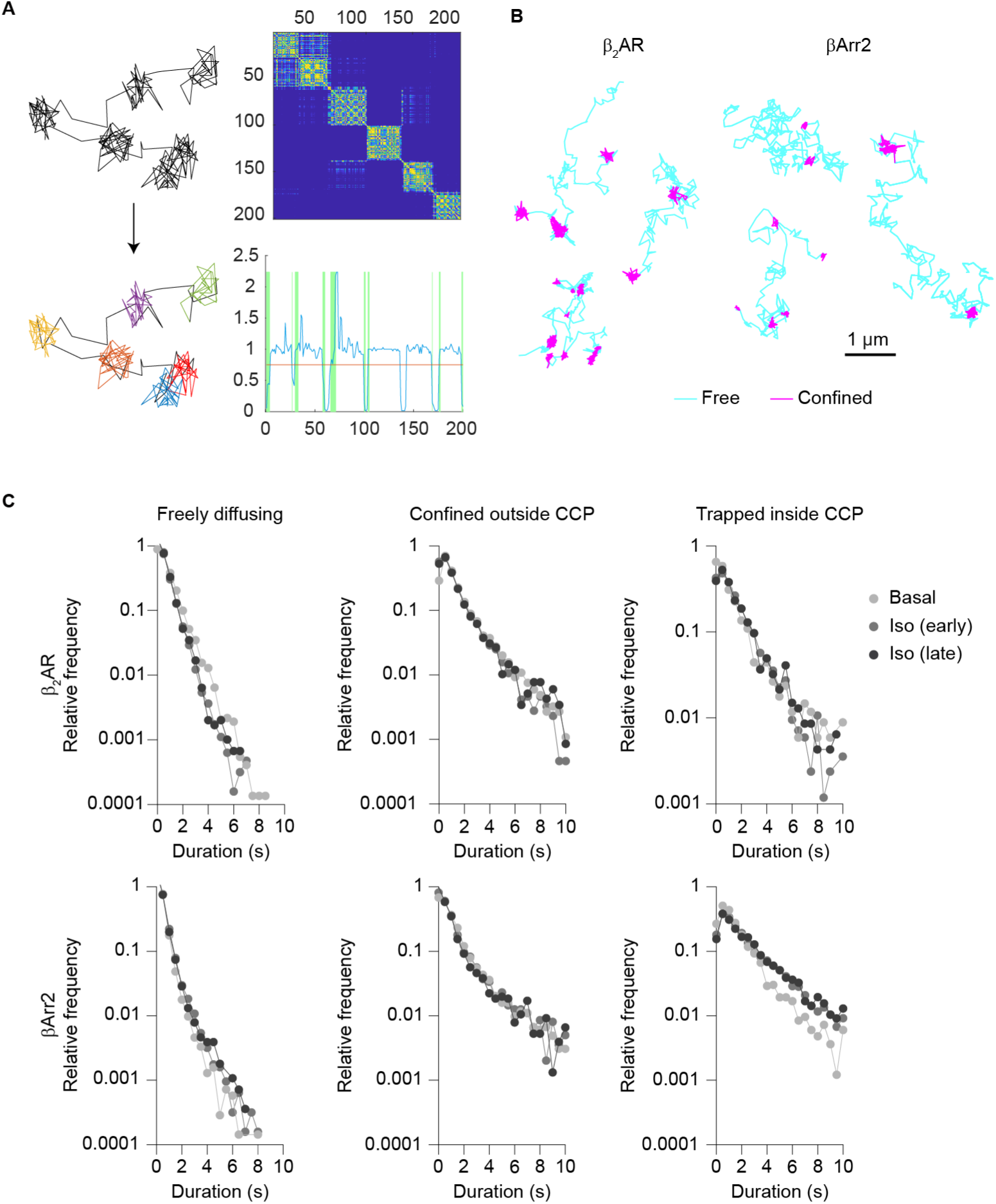
Spatial Confinement Analysis, Related to Figure 1. (A) Detection of spatially confined trajectory segments. Left, simulated trajectory alternating phases of free and confined diffusion (top) and result of segmentation (bottom). Segments characterized by free diffusion are shown in black, whereas confined ones are highlighted in different colors. Right, corresponding recurrence matrix (top) and discriminator (bottom) used to identify confined trajectory segments. (B) Application of the analysis to β_2_AR and βArr2 trajectories. Shown are representative trajectories alternating between phases of free diffusion and spatial confinement. (C) Distributions of the durations of β_2_AR (top) and βArr2 (bottom) trajectory segments characterized by free diffusion, confinement outside CCPs, and confinement (i.e., trapping) inside CCPs. Shown are the results before (basal) and after stimulation with isoproterenol (10 iJM). lso, isoproterenol. Early, 2-7 min. Late, 8-15 min.

**Figure S4.**
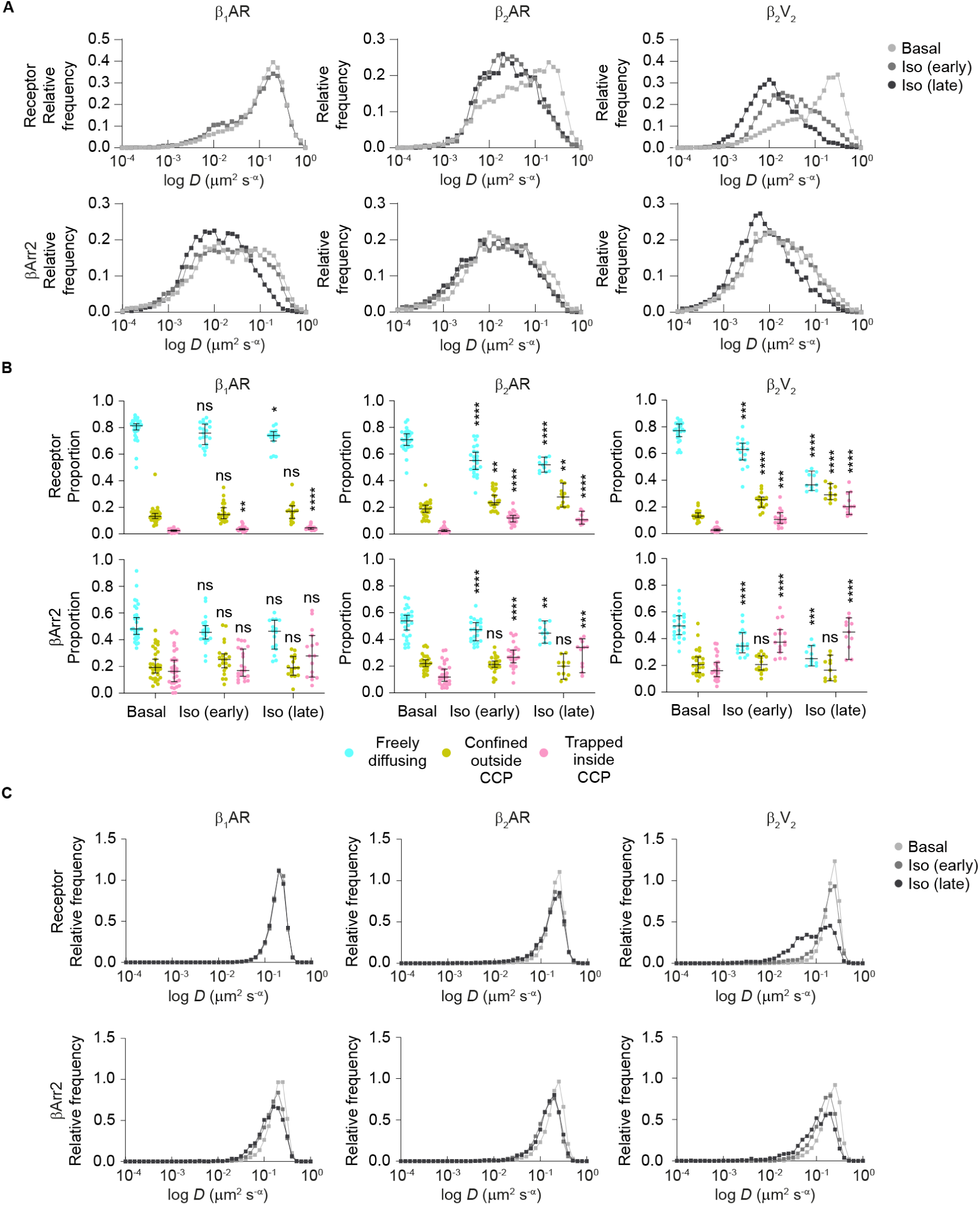
Additional Results of TAMSD and Spatial Confinement Analyses, Related to Figure 2. Shown are results obtained in cells expressing βArr2-Halo and either SNAP-β_1_AR, −β_2_AR, or −β_2_V_2_ before (basal) and after isoproterenol (10 μM) stimulation. β_2_AR is included for comparison. (A) Distributions of diffusion coefficient (*D*) values estimated for receptor (top) and βArr2 (bottom) molecules based on TAMSD analysis of entire trajectories. (B) Lateral mobility of receptor (top) and βArr2 (bottom) molecules. Three diffusivity states were identified based on the spatial confinement analysis and location at CCPs. Results are displayed as in Figure 1G. (C) Distributions of diffusion coefficients obtained from TAMSD analysis of free trajectory portions. Iso, isoproterenol. Early, 2-7 min. Late, 8-15 min. Data in B are median ± 95% confidence interval. n = 31, 21, 16, 28, 23, 11, 24, 16, 11 cells for β_1_AR basal, β_1_AR Iso (early), β_1_AR Iso (late), β_2_AR basal, β_2_AR Iso (early), β_2_AR Iso (late), β_2_V_2_ basal, β_2_V_2_ Iso (early), β_2_V_2_ Iso (late), respectively. Results in B are statistically significant by Kruskal Wallis test. *P < 0.05, **P < 0.01, ***P < 0.001, ****P < 0.0001 versus basal by t-test with Bonferroni correction.

**Figure S5.**
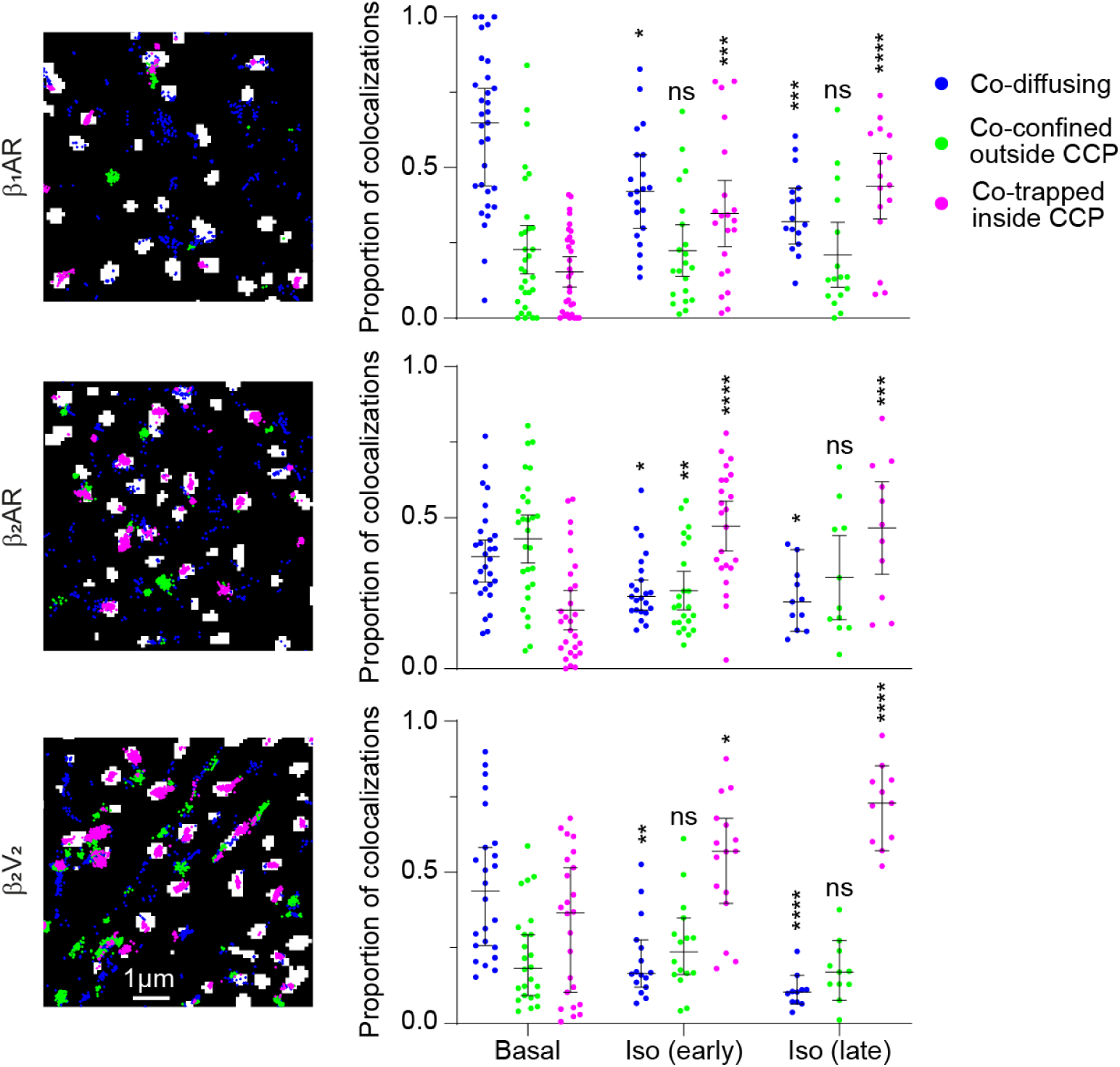
Additional Results of Spatial Analysis of Receptor–βArr2 Single-Molecule Colocalization Events, Related to Figure 2. Shown are the results obtained in cells expressing βArr2-Halo and either SNAP-β_1_AR, −β_2_AR, or −β_2_V_2_ before (basal) and after isoproterenol (10 μM) stimulation. Results are shown as in Figure 1K. β_2_AR is included for comparison. Iso, isoproterenol. Early, 2-7 min. Late, 8-15 min. Data are median ± 95% confidence interval. Results are statistically significant by Kruskal Wallis test. *P < 0.05, **P < 0.01, ***P < 0.001, ****P < 0.0001 versus basal by t-test with Bonferroni correction. n = 31, 21, 16, 28, 23, 11, 24, 16, 11 cells for β_1_AR basal, β_1_AR Iso (early), β_1_AR Iso (late), β_2_AR basal, β_2_AR Iso (early), β_2_AR Iso (late), β_2_V_2_ basal, β_2_V_2_ Iso (early), β_2_V_2_ Iso (late), respectively. Images are representative of a minimum of 3 independent experiments.

**Figure S6.**
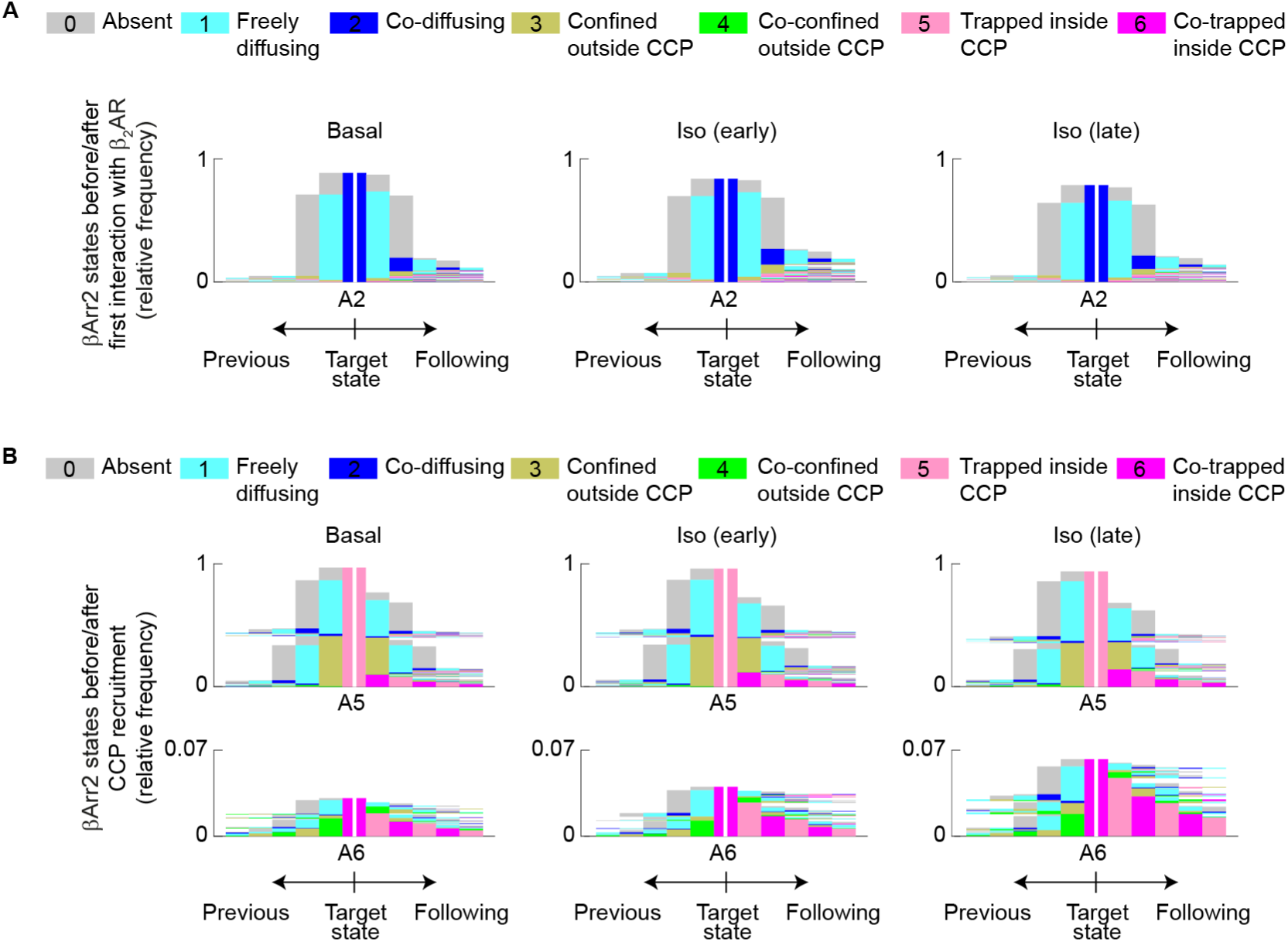
History Plots, Related to Figure 3. (A) History plots summarizing the relative frequencies of all observed sequences of βArr2 states preceding and following the state corresponding to β_2_AR–βArr2 co-diffusion (A2). (B) History plots summarizing the βArr2 states preceding and following the two states (A5, A6) corresponding to βArr2 trapped inside a CCP. Data before and after each target state in A and B are sorted independently to facilitate visualization. Iso, isoproterenol. Early, 2-7 min. Late, 8-15 min. n = 88,851/51,147, 52,658/58,254, 27,741/22,769 transitions for β_2_AR/βArr2 basal, β_2_AR/βArr2 Iso (early), β_2_AR/βArr2 Iso (late), respectively.

**Figure S7.**
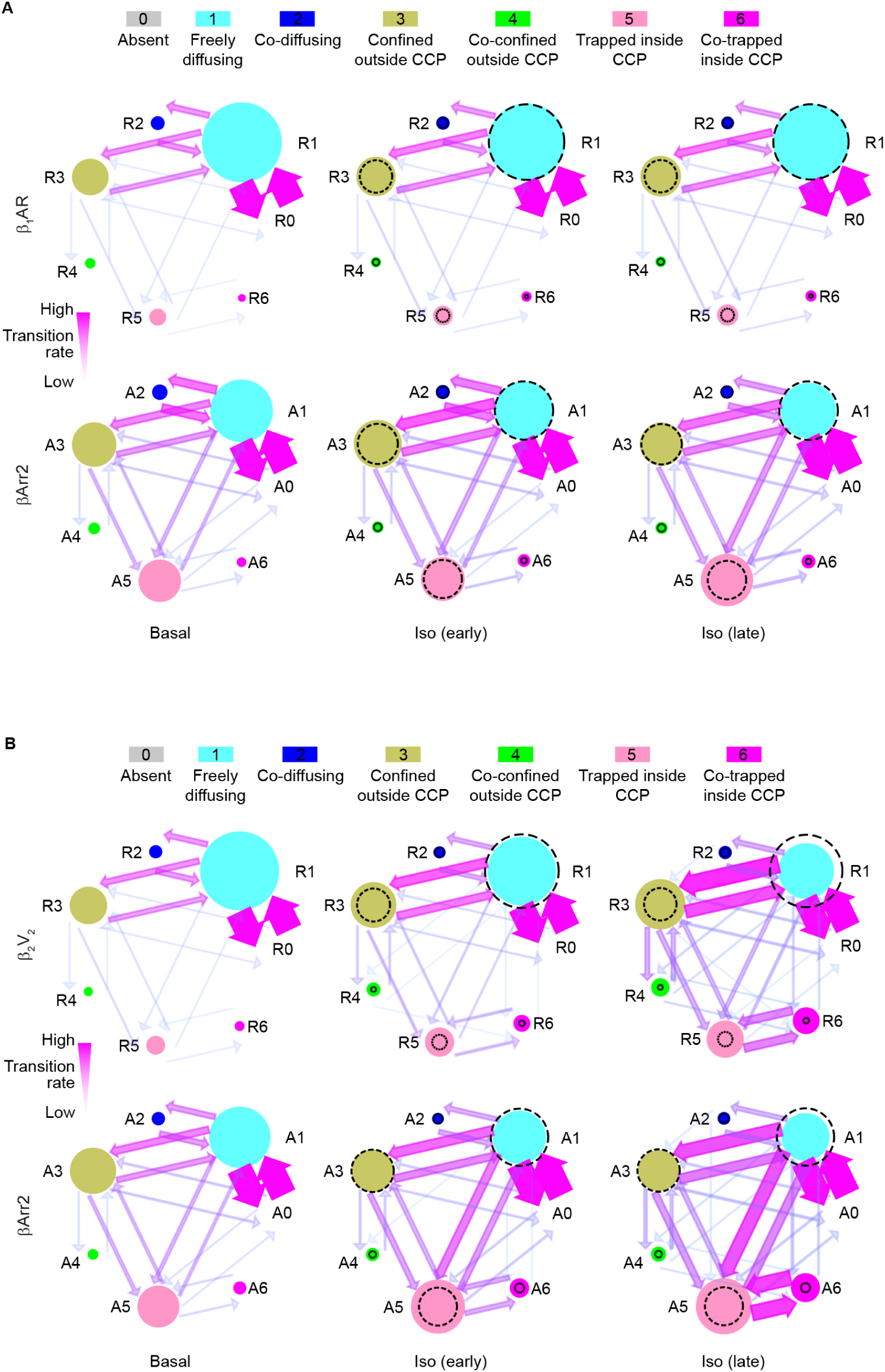
Sequence of Events in β_1_AR–βArr2 and β_2_V_2_–βArr2 Interactions, Related to Figure 3. (A, B) Results of Markov chain analysis applied to β_1_AR–βArr2 (A) and β_2_V_2_–βArr2 (B). Data are shown as in Figure 3A. Iso, isoproterenol. Early, 2-7 min. Late, 8-15 min. n = 125,983/65,731, 79,226/47,650, 67,672/38,993 transitions (A) for J3,AR/J3Arr2 basal, J3,AR/J3Arr2 lso (early), J3,AR/J3Arr2 lso (late), and 82,036/44,390, 49,470/40,762, 32,797/29,085 transitions (B) for J3N2/J3Arr2 basal, J3N2/J3Arr2 lso (early), J3N2/J3Arr2 lso (late), respectively.

**Figure S8.**
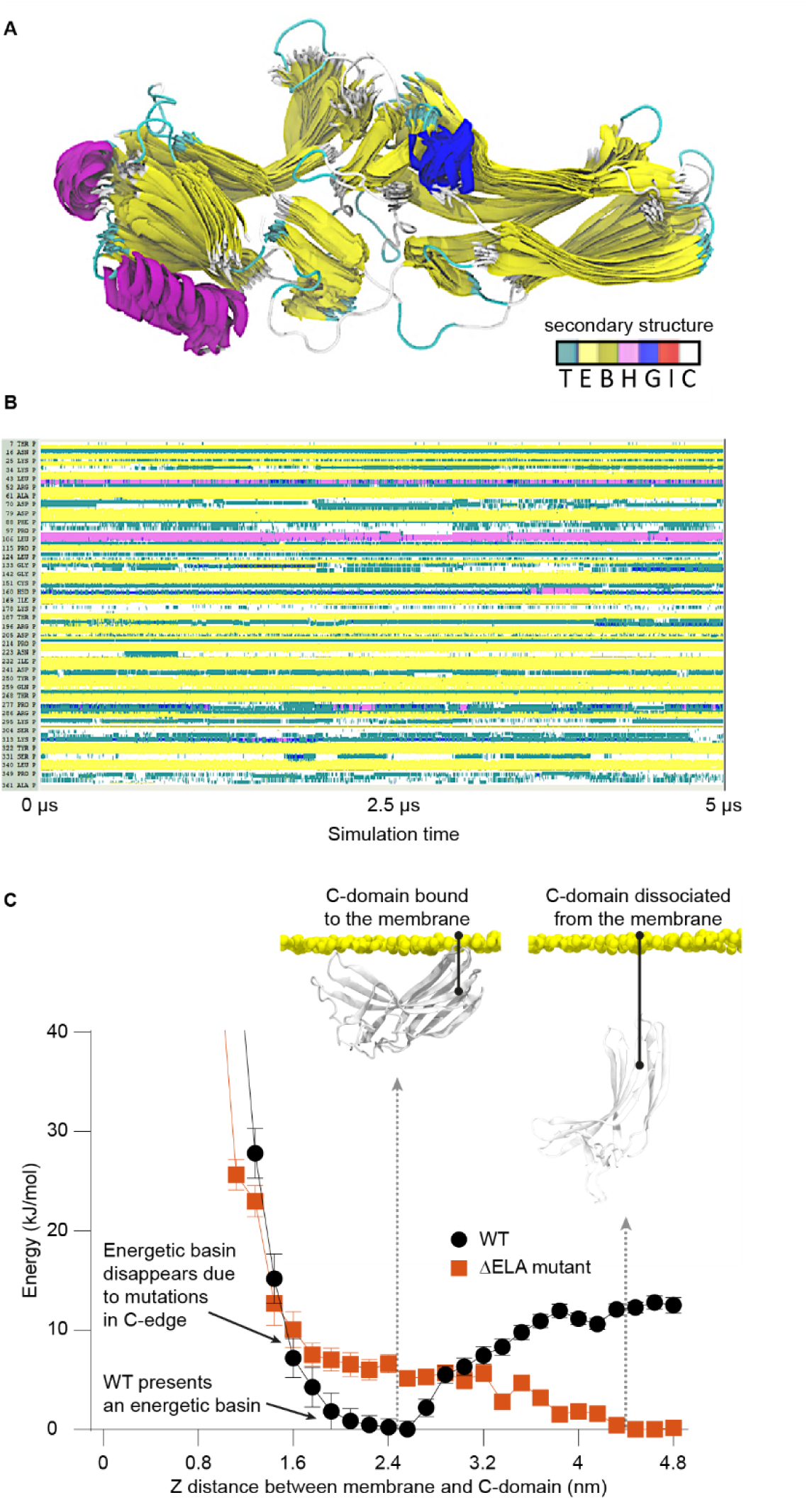
MD Simulations of the ΔELA Mutant, Related to Figure 5. (A) Stability of the ΔELA mutant evaluated by MD simulations (accumulated time 5 x 1 μs). Shown is the overlay of βArr2 snapshots separated by 120 ns intervals. For simplicity, only the evolution of elements with defined secondary structure is shown. (B) Evolution of the secondary structure of βArr2 during the simulations. The secondary structure was calculated using the timeline tool integrated in the VMD package. T, turn. E, extended configuration. B, isolated bridge. H, α-helix. G, 3_10_ helix. I, π-helix. C, coil. (C) Well-tempered metadynamics simulations comparing the t.ELA mutant and wild-type 0fVT) J3Arr2. Shown are the free energy landscapes as the proteins are pulled out of the membrane using as collective variable the distance between the C-domain and the plasma membrane.

**Figure S9.**
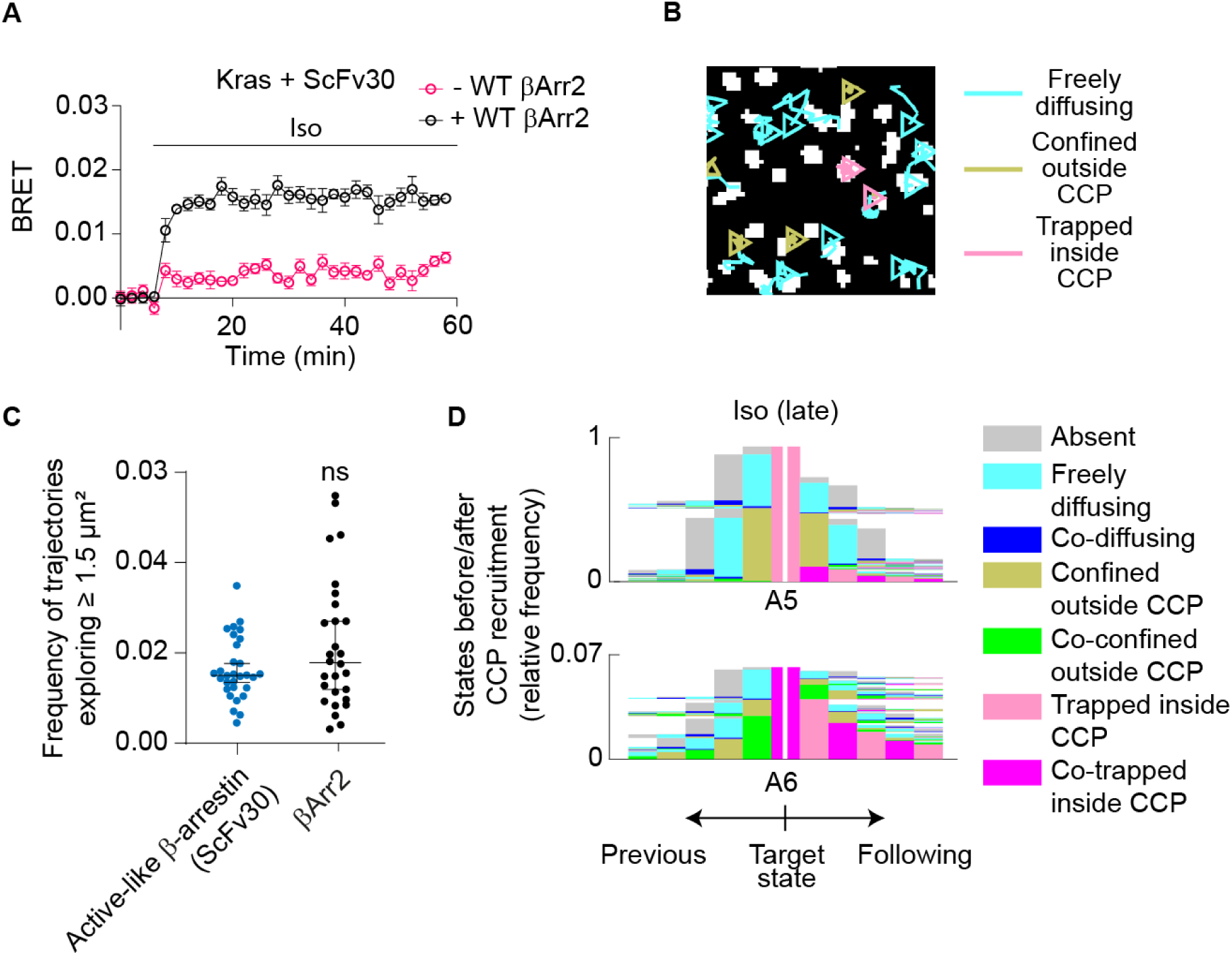
Additional Results with Scfv30, Related to Figure 6. (A) Kinetics of ScFv30 recruitment to the plasma membrane in βArr1/2 CRISPR-Cas9 knockout HEK cells with/without βArr2 re-expression upon isoproterenol (10 μM) stimulation. (B) Diffusivity states of active-like β-arrestin on the plasma membrane as revealed by single-molecule visualization with ScFv30. Shown are representative trajectories in cells stimulated with isoproterenol (10 μM) for 8-15 min (late). (C) Propensity of active-like β-arrestin detected by ScFv30 to explore the plasma membrane. Total βArr2 is given for comparison. Shown are the relative frequencies of molecules exploring ≥ 1.5 μm^2^ in unstimulated cells. (D) History plots summarizing the states of active-like β-arrestin (ScFv30) preceding and following the two states (A5, A6) corresponding to trapping inside a CCP. Data are shown as in Figure S6. Data are mean ± s.e.m. (A) and median ± 95% confidence interval (C). n = 3 biological replicates (A), 30, 28 cells for ScFv30, total J3Arr2 (C), and 99,428 transitions for ScFv30 lso (late) (D). ns, statistically not significant by Kruskal Wallis test (C). The image in B is representative of 3 independent experiments.

## METHODS

### Resource availability

#### Lead contact

Further information and requests for resources and reagents should be directed to and will be fulfilled by the lead contact, Davide Calebiro.

#### Materials availability

Plasmids generated in this study are available from the lead contact. This study did not generate new unique reagents.

#### Data and code availability

MATLAB scripts will be uploaded to an online platform and made available to the public upon manuscript acceptance.

### Experimental model and subject details

#### Materials

Cell culture reagents, Lipofectamine 2000 (cat. no. 11668019) and TetraSpeck fluorescent beads (cat. no. T7279) were purchased from Thermo Fisher Scientific. Isoproterenol (cat. no. 1747) was from Tocris Bioscience. The fluorescent benzylguanine derivative SNAP-Surface Alexa Fluor 647 (cat. no. S9136S) was from New England Biolabs. HaloTag Janelia Fluor 549 (cat. no. GA111A), HaloTag R110Direct (cat. no. G322A) and furimazine (cat. no. N1120) were from Promega. Ultraclean glass coverslips were obtained as previously described (Calebiro et al., 2013).

#### Cell culture and transfection

Chinese hamster ovary K1 (CHO-K1) cells (ATCC) were cultured in phenol red-free Dulbecco’s modified Eagle’s medium (DMEM)/F12 (cat. no. 11039-021), supplemented with 10% FBS (cat. no. 10500-064), 100 U/ml penicillin and 0.1 mg/ml streptomycin (cat. no. 15140-122) at 37 °C, 5% CO_2_. Cells were seeded onto ultraclean 25-mm round glass coverslips at a density of 3 x 10^5^ cells per well. On the next day, they were transfected using Lipofectamine 2000, following the manufacturer’s protocol. Cells were labeled and imaged by single-molecule microscopy 3.5-4 hours after transfection to obtain low physiological expression levels (Calebiro et al., 2013; Sungkaworn et al., 2017).

For BRET experiments, human embryonic kidney 293 (HEK293) (ATCC) or HEK293 βArr1/2 CRISPR KO cells (kindly provided by Asuka Inoue) (O’Hayre et al., 2017; Schrage et al., 2015) were cultured in DMEM (cat. no. 41966-029), supplemented with 10% FBS, 100 U/ml penicillin and 0.1 mg/ml streptomycin at 37 °C, 5% CO_2_. Cells were transfected with Lipofectamine 2000 following the manufacturer’s protocol. After 24 hours, they were resuspended in FluoroBrite phenol red-free DMEM medium (cat. no. A18967-01) supplemented with 4 mM L-glutamine and 5% FBS and plated into poly-D-lysine-coated 96-well white polystyrene Nunc microplates (Sigma) at a density of 1×10^5^ cells/well. Forty-eight hours post transfection, the medium was replaced with HBSS (cat. no. 14025-050) containing 100 nM Halo R110 ligand and incubated for 1 hour at 37 °C for labeling, followed by addition of 10 μM furimazine.

Cells were routinely tested for mycoplasma contamination.

### Method details

#### Molecular biology

Plasmids encoding N-terminally SNAP-tagged human β_1_AR (SNAP-β_1_AR), β_2_AR (SNAP-β_2_AR) and CD86 (SNAP-CD86) were previously generated and verified to be functional (Calebiro et al., 2013). A plasmid encoding N-terminally SNAP-tagged human β_2_AR carrying the vasopressin V_2_ receptor C-tail (SNAP-β_2_V_2_) was generated by replacing the C-tail in the SNAP-β_2_AR construct with that of V_2_ (ARGRTPPSLGPQDESCTTASSSLAKDTSS). Plasmids encoding N-terminally SNAP-tagged human β_2_AR with a deletion in the third intracellular loop (SNAP-β_2_AR ΔICL3) or lacking the entire C-tail (SNAP-β_2_AR ΔC-tail) were generated by PCR using standard procedures. A plasmid encoding C-terminally Halo-tagged bovine βArr2 (βArr2-Halo) was generated by replacing CFP with the Halo tag in a previously described βArr2-CFP construct (Nuber et al., 2016). A plasmid encoding C-terminally Halo-tagged bovine βArr2 carrying mutations interfering with binding to both clathrin (L373A/I374A/F376A) and AP2 (R393A/R395A) (Laporte et al., 2000; Laporte et al., 1999) (βArr2 ΔCCPAP2-Halo) was generated by PCR mutagenesis. Similar procedures were used to generate plasmids encoding C-terminally Halo-tagged bovine βArr2 with a deletion of the finger loop (YGREDLDVLGLSFRK) (βArr2 ΔFLR-Halo) (Cahill et al., 2017) or carrying mutations (K233Q/R237Q/K251Q) that interfere with PIP2 binding (βArr2 ΔPIP2-Halo) (Gaidarov et al., 1999; Milano et al., 2002). An additional βArr2-Halo construct carrying a panel of mutations (R189Q/F191E/L192S/M193G/T226S/K227E/T228S/K230Q/K231E/K233Q/R237E/K251Q/K 325Q/K327Q/V329S/V330D/R332E) designed to prevent plasma membrane interactions (βArr2 ΔELA-Halo) was generated by PCR mutagenesis of the βArr2 ΔPIP2-Halo construct, followed by Gibson assembly. A plasmid encoding human β_2_AR containing NanoLuciferase (Nluc) fused to its C-terminus (β_2_AR-Nluc) was previously described (Kilpatrick et al., 2019). Plasmids encoding β_1_AR, β_2_V_2_, β_2_AR ΔICL3 and β_2_AR ΔC-tail with Nluc fused to their C-termini were cloned from the β_2_AR-Nluc construct by replacing the β_2_AR coding sequence with those of the corresponding receptor constructs. A plasmid encoding K-Ras with Nluc fused to its N-terminus was kindly provided by Kevin Pfleger (White et al., 2017). A plasmid encoding N-terminally GFP-tagged clathrin light chain (GFP-CCP) was kindly provided by Emanuele Cocucci and Tom Kirchhausen (Cocucci et al., 2012). A plasmid encoding N-terminally CFP-tagged clathrin light chain (CFP-CCP) was cloned by replacing GFP in GFP-CCP using PCR and Gibson assembly. A plasmid encoding Lifeact-YFP was generated by replacing GFP with YFP in a previously described Lifeact-GFP construct (kindly provided by Antje Gohla) (Sungkaworn et al., 2017). A plasmid encoding N-terminally Nluc-tagged clathrin light chain was obtained by gene synthesis (Twist Bioscience). A plasmid encoding C-terminally Halo-tagged ScFv30 (ScFv30-Halo) was generated by replacing the YFP sequence with Halo in a previously described ScFv30-YFP construct (Min et al., 2020; Pandey et al., 2021).

### BRET measurements

BRET measurements were performed at 37 °C using a PHERAstar Microplate Reader (BMG Labtech) with a dual-luminescence readout BRET1 plus filter (460-490 nm band-pass, 520-550 nm long-pass). Following 4 baseline measurements, the cells were treated with vehicle or 10 μM isoproterenol and measured for an additional hour. BRET acceptor/donor ratios were calculated separately for each well. The results were normalized to the baseline values and those obtained with vehicle. Measurements were performed in triplicate readouts.

### Live cell protein labeling for single-molecule microscopy

Cells were labeled with a combination of 1 μM SNAP-Surface Alexa Fluor 647 (AF647, cell impermeable) and 1 μM HaloTag Janelia 549 (JF549, cell permeable) in complete culture medium for 20 min at 37 °C. These concentrations were selected based on titration experiments to obtain saturation labeling of both SNAP- and Halo-tagged proteins (**Figure S1C**), which results in ~90% and ~70% labeling efficiencies for extracellular and intracellular tags as determined by single-molecule microscopy (Calebiro et al., 2013). Cells were then washed three times with complete culture medium, allowing 5 min incubation between washes. Non-specific labeling was <1% (**Figure S1D**).

### Single-molecule microscopy

Single-molecule microscopy experiments were performed using total internal reflection fluorescence (TIRF) illumination on a custom system (assembled by CAIRN Research) based on an Eclipse Ti2 microscope (Nikon, Japan) equipped with a 100x oil-immersion objective (SR HP APO TIRF NA 1.49, Nikon), 405, 488, 561, and 637 nm diode lasers (Coherent, Obis), an iLas2 TIRF illuminator (Gataca Systems), quadruple band excitation and dichroic filters, a quadruple beam splitter, 1.5x tube lens, four EMCCD cameras (iXon Ultra 897, Andor), hardware focus stabilization, and a temperature-controlled enclosure. The sample and objective were maintained at 37 °C throughout the experiments. Coverslips were mounted in a microscopy chamber filled with Hank’s balanced salt solution (HBSS) supplemented with 10 mM HEPES (cat. no. H0887), pH 7.5. A reduced oxygen environment (2-4% O_2_) was provided in the imaging chamber to decrease photobleaching without increasing cytotoxicity using a mixture of nitrogen and air and a home-built gas mixing and humidifying system as previously described (Tsunoyama et al., 2018). The oxygen concentration in the imaging solution was measured in real-time using a needle-type oxygen sensor connected to an OXY-1 microsensor (OXY-1 ST PreSens, Germany). Multi-color single-molecule image sequences were acquired simultaneously on the four synchronized EMCCDs at a rate of one image every 33 ms.

#### Single-particle tracking

Automated single-particle detection and tracking were performed with the u-track software (Jaqaman et al., 2008) and the obtained trajectories were further analyzed using custom algorithms in MATLAB environment as previously described (Sungkaworn et al., 2017). Image sequences from different channels were registered against each other using a linear piecewise transformation, based on reference points taken with multi-color fluorescent beads (100 nm, TetraSpeck) (Sungkaworn et al., 2017). The inter-channel localization precision after coordinate registration was ~20 nm.

#### β-arrestin purification and labeling

A minimal cysteine mutant of bovine β-arrestin 1 carrying an N-terminal GST-tag separated by a thrombin-cleavage site and cloned in the pGEX4T3 expression vector was used (Kumari et al., 2016). Two additional amino acids, Ala-Cys, were introduced at the C-terminus to allow site-specific labeling and the construct was confirmed by DNA sequencing. GST-β-arrestin was expressed in *E. coli* BL21(DE3) cells (New England Biolabs, cat. no. C2527I). A starter culture grown in LB broth (Sigma, cat. no. L3022) supplemented with 100 μg/ml ampicillin (Sigma, cat. no. A9518) at 37 °C to an A600 of 0.6 was used to inoculate 2 liters of Terrific Broth (Fisher Scientific, cat. no. 22711022) supplemented with 100 μg/ml ampicillin, which was also grown at 37 °C. When A600 reached 0.6-0.8, the cultures were equilibrated to 18 °C, and expression was induced with 50 μM Isopropyl β-D-1-thiogalactopyranoside (Sigma, cat. no. I6758) for 16 h. The cells were harvested in PBS, resuspended in 180 ml of cold lysis buffer (25 mM Tris-HCl pH 8.5, 150 mM NaCl, 1 mM Phenylmethanesulfonyl Fluoride, 2 mM Benzamidine hydrochloride, 1 mM EDTA, 5% glycerol, 2 mM DTT) with the addition of 10 μl Benzonase nuclease (Sigma, cat. no. E1014-25KU), lysed by sonication and cleared by ultracentrifugation at 100,000 g at 4 °C for 40 min. All following steps were done at 4 °C unless otherwise stated. The cleared lysate was filtered through a 0.22-μm syringe filter, applied to 20 ml of Glutathione Sepharose 4B (Cytiva, cat. no. GE17-0756-01) resin, pre-equilibrated in wash buffer (50 mM Tris-HCl pH 8.5, 150 mM NaCl) and incubated overnight under gentle rotation. The Sepharose suspension was spun at 1,000 g for 15 min, the supernatant decanted and the resin transferred to a glass chromatography column filled with wash buffer. The resin was washed with 20 column volumes (CV) of high-salt wash buffer (50 mM Tris-HCl pH 8.5, 1 M NaCl), and 10 CV of wash buffer. GST-β-arrestin was eluted in 1 CV fractions of elution buffer (50 mM Tris-HCl pH 8.5, 150 mM NaCl, 2 mM DTT, 20 mM Glutathione), the protein content was estimated by Pierce BCA Protein Assay (Fisher Scientific, cat. no. 10741395), and the fractions containing proteins pooled and concentrated to 5 ml. The buffer was adjusted to 350 mM NaCl and 0.02% n-dodecyl β-D-maltoside (DDM), thrombin protease (250 U, Sigma, cat. no. T7572-250UN) was added, and cleavage was allowed for 2 h at room temperature before it was stopped by the addition of 2 mM benzamidine hydrochloride (Sigma, cat. no. 434760-5G). The solution was concentrated to 600 μl and β-arrestin was isolated on a Superdex 200 Increase 10/300 GL column (Cytiva) equilibrated in 25 mM Tris-HCl pH 8.5, 350 mM NaCl, 0.02% DDM at room temperature. Peak fractions were pooled, concentrated to 250 μl, and the pH was adjusted by the addition of 40 mM Tris-HCl pH 6.8. Disulphide bridges were reduced by the addition of 0.8 μM Tris(2-carboxyethyl)phosphine hydrochloride (Sigma, cat. no. C4706). β-arrestin was labeled by incubation with 0.8 μM Alexa Fluor™ 647 C2 Maleimide (Invitrogen, cat. no. A20347) for 2 h at room temperature in the dark, followed by polishing on a Superdex 200 Increase 10/300 GL column as described above. Peak fractions were pooled and aliquots flash-frozen in liquid nitrogen and stored at −80 °C.

#### Giant unilamellar vesicle preparation

Giant unilamellar vesicles (GUVs) were obtained by electroformation. A total of 7 μl of di-oleoyl-phosphatidylcholine lipid solution (10 mg/ml in chloroform) was added to indium tin oxide-coated glass slides to form a lipid film on the conductive surface. Once the lipid films were dry, a chamber with 0.3-mm gap was assembled and filled with a 200 mM sucrose solution. The assembled chamber was then placed at 50°C and the following conditions were applied for GUV electroformation: 11 Hz, 1V alternating electric current for 2 h.

#### Experiments with supported lipid bilayers

Custom imaging chambers were assembled as previously described (Jain et al., 2012). Briefly, 0.75-mm diameter inlet/outlet holes were drilled in glass microscopy slides at a distance of 3-4 mm from the edges to create a flow channel. Coverslips (VWR, 24 x 40 mm) were cleaned by sequential sonication in chloroform and 5 M NaOH solution, rinsed in distilled water and allowed to dry. To assemble the flow chambers, double-sided Scotch tape was sandwiched between a slide and a coverslip and the edges were sealed with an epoxy glue (5-Minute Epoxy, Thorlabs), resulting in the formation of flow channels connected to the inlet/outlet holes. The chambers were rinsed with 200 μl PBS. A total of 20 μl of the GUV suspension was mixed with 0.2 μl a lipophilic dye solution (5 μM 3,3’-Dioctadecyloxacarbocyanine perchlorate in N,N-Dimethylformamide) and loaded into the chambers, followed by incubation for 1 h at room temperature in the dark to allow for the GUVs to break through osmotic shock and form lipid bilayers. Afterwards, the chambers were rinsed with 200 μl PBS, followed by 200 μl PBS containing 0.1% bovine serum albumin (BSA). Purified β-arrestin was suspended to a concentration of 1 nM in an oxygen-scavenging buffer (10 mM Tris-HCl pH 7.4, 50 mM NaCl, 0.1% BSA, 1% D-glucose, 2 mM Trolox, 25 U/ml glucose oxidase, 250 U/ml catalase) and loaded into a chamber, which was immediately sealed and imaged by single-molecule microscopy as described above for live cells.

#### Single-molecule interaction analysis and estimation of *k*_on_/*k*_off_ values

The frequency and duration of receptor–βArr2 interactions were estimated using our previously described method based on deconvolution of the distribution of single-molecule colocalization times with the one expected for random colocalizations (Sungkaworn et al., 2017). The distribution for random colocalizations was estimated in cells co-transfected with βArr2-Halo and SNAP-tagged CD86, a non-related membrane protein that does not interact with βArr2 and has diffusion characteristics comparable to those of the investigated receptors. Single-molecule interactions and microscopic *k*_on_/*k*_off_ values were estimated as previously described (Sungkaworn et al., 2017). Briefly, for each particle in channel 1 at each frame, all particles in channel 2 falling within a defined search radius (*R*_0_ = 150 nm) were identified as colocalizing. For each colocalization event, the starting and terminating frame was obtained. A Monte Carlo approach was used to link colocalization events that were prematurely terminated due to uncertainty in the assignment of trajectory segments after a splitting event. Data obtained in cells expressing SNAP-CD86 labeled SNAP-AF647, βArr2-Halo labeled Halo-JF549 and unlabeled wild-type β_2_AR were used to estimate the frequency and duration of random colocalization.

The distributions of true interaction times were estimated by deconvolving the observed distributions of colocalization times with that obtained for the non-interacting control pair (SNAP-CD86 and βArr2-Halo).

To estimate dissociation rate constant (*k*_off_) values, normalized relaxation curves obtained from the distributions of true interaction times were fitted to an exponential decay function:

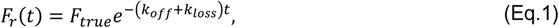

where *F*_r_(*t*) is the fraction of surviving interactions at time *t*, *F*_true_ (≤1) is the fraction of true interactions (over total interactions) at *t* = 0, and *k*_loss_ is a correction factor accounting for premature termination of the interactions due to photobleaching or potential particle loss at detection or tracking. *k*_loss_ was estimated based on simulated image sequences of randomly diffusing, non-dissociating particles with characteristics (particle densities, point spread functions, diffusion coefficients, background levels, signal-to-noise ratios, fluorophore bleaching rates) matching the experimental ones.

Association rate constant (*k*_on_) values are related to *F*_true_ and the rate of new co-localizations per unit of area *d*[*C*]_ρ_/*dt* by the following equation:

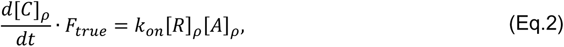

where []_ρ_ denotes density and [*R*]_ρ_ and [*A*]_ρ_ are the densities of free receptor and β-arrestin molecules, which can in turn be derived by subtracting the estimated density of receptor–β-arrestin complexes [RA]ρ) from the total densities measured in channels 1 [*C*ℎ1]*ρ* and 2 [*C*ℎ2]*ρ*, respectively.

Finally, [*RA*]_ρ_ can be deduced based on the balance between association and dissociation rates at equilibrium using the formula:

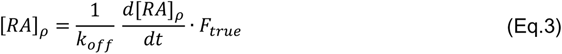

These relationships allowed us to estimate *k_on_* from measured observables using to the following formula:

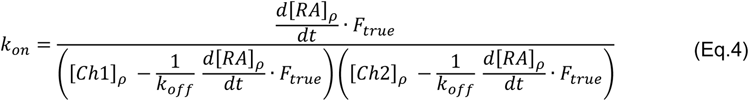

*k*_on_ was estimated separately at each frame and the given values were obtained by averaging over the analyzed frames.

#### TAMSD analysis

The time-averaged mean squared displacement (*TAMSD*) (Kepten et al., 2015; Lanoiselée et al., 2018) of individual trajectories was computed as previously described (Sungkaworn et al., 2017). *TAMSD* data were fitted to the equation describing the ensemble averaged *TAMSD* for an ergodic process:

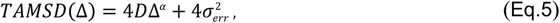

where Δ indicates lag time, *α* is the anomalous diffusion exponent and *σ*_err_ is a constant offset accounting for localization error. Only trajectories lasting at least 100 frames were analyzed.

Data corresponding to the first 10 lag time points were used for the fitting.

For the analysis of sub-trajectories, all trajectory segments lasting at least 50 frames were included. Segments characterized by free diffusion were fitted using Eq. 5. Segments characterized by confinement/trapping were fitted using the equation describing the diffusion of a molecule inside a harmonic potential (Cherstvy et al., 2018):

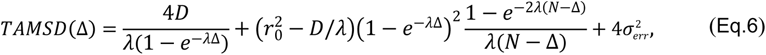

Where 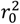 is the squared distance from the center of the trap at the beginning of the diffusive process and *λ* is the inverse of the mean reversion time. From this, the approximate confinement/trap diameter (*d*) was deduced as 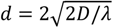.

#### CCP detection

CCP detection was performed by applying a frame-by-frame binary mask to GFP-CCP image sequences. Image sequences were pre-processed to remove local background and enhance contrast using a bottom and top-hat filter with a disk-shaped structuring element of 11-pixel diameter (’imtophat’ and ‘imbothat’ functions in MATLAB). A Kalman filter with low gain (0.1) was applied to reduce noise. The image sequences were then deconvolved with the theoretical PSF of the system using the Lucy-Richardson algorithm (Lucy, 1974; Richardson, 1972). Binary masks corresponding to CCPs were obtained by thresholding the image sequences with a value corresponding to the mean plus 1 standard deviation of each frame. Only pixels persisting at least 45 consecutive frames (~1.5 s) were included in the final CCP masks.

#### Spatial confinement analysis

A spatial confinement analysis was used to identify trajectory segments characterized by confinement/trapping, using our recently described algorithm (Lanoiselée et al., 2021). Briefly, for each trajectory, we computed a recurrence matrix (M*ij*) containing information about the distance between each pair of points in the trajectory:

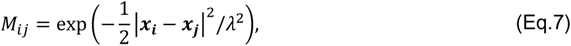

where *i* and *j* run over each step of the trajectory, ***x*** is the position of the particle, and λ is the test length scale of the analysis. *M_ij_* approaches 1 if the difference between ***x_i_*** and ***x_j_*** is smaller than λ. To minimize the effects of localization error and possible misdetected outliers, *M_ij_* was smoothened by local averaging and thresholded to obtain a binary matrix (*B*), where *B_ij_* = 1 if M*_ij_* > *e*^−1^ or zero otherwise. Trapped portions of the trajectory appear in *B* as square blocks of ones along its diagonal. To detect these blocks, three quantities were calculated, i.e. block time (*t*_|_ (*n*)), neighbouring time (*t*_┴_ (*n*)) and persistence time (*t*_‖_ (*n*)), from which we computed the invariant quantity *v*(*n*):

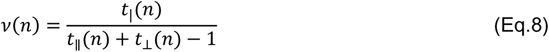

In the idealized case of a perfect squared box of ones it is easy to verify that *v*(*n*) = 1. In practice, blocks are never perfect squares, and we use a cut-off of *v_n_* = ¾ to identify blocks corresponding to potential confined trajectory segments. A statistical test based on the probability of detecting a larger block by chance for a particle with 2D fractional Brownian motion and P value = 0.05 is then applied to decide if the detected block is a confinement event.

#### Markov chain analysis

A Markov chain analysis was used to estimate the relative occupancies and forward transition probabilities. Each molecule at each frame was labeled with four binary numbers describing the presence/absence of spatial confinement, CCP localization, co-localization with either a β-arrestin molecule (in case of a receptor) or a receptor (in case of β-arrestin), and confinement of the colocalizing partner. Considering all possible combinations and discarding the physically irrelevant ones, such as freely diffusing molecules on a CCP or colocalization between a freely diffusing and a confined molecule, we obtained a set of 6 unique states plus a dummy state corresponding to molecules before or after their detection on the plasma membrane to account for movements between the cytosol and the plasma membrane as well as disappearance due to fluorophore photobleaching (**Table S2**). Forward Markov chains were built by gathering state information from all trajectories. Only states lasting at least 10 frames (0.33 s) and their transitions were considered.

#### History plots

Plots displaying all the observed sequences of events before and after a given target state were generated in several steps. First, we collected all the trajectories that contained the target state. Then, for each trajectory, we separately gathered the *h* states preceding and following the target state. The information was used to build two separate tree graphs (past and future), where the branches represent all observed sequences of events. Taking advantage of graph theory, we assigned a branch number to each history in either graph using the following formula:

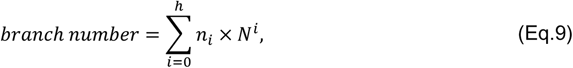

where *i* runs over all states in the sequence (from the target state *i* = 0 to the farthest state from present to be analyzed for *i* = *h*) with *n_i_* being the state number of the state *i* and *N* the number of distinct states (here *N* = 7).

Finally, histories were sorted according to their branch number. Graphic representations were obtained by stacking all histories, each represented by a thin horizontal line, color coded according to the contained states.

#### Super-resolution radial fluctuation imaging

Super-resolved Lifeact images were generated from image sequences using the super-resolution radial fluctuations (SRRF) algorithm implemented in the NanoJ-SRRF ImageJ plugin (Gustafsson et al., 2016). In overlaid images, the coordinates of single-molecule trajectories and CCPs binary masks were rescaled to match the higher resolution of SRRF images.

#### Molecular dynamics

The initial conformation of βArr2 was modelled with MODELLER (Fiser et al., 2000; Sali and Blundell, 1993) using the structure of βArr1 in complex with the muscarinic M2 receptor (PDB code: 6U1N) (Staus et al., 2020). This template was selected as the C-edge is resolved in a conformation that exposes hydrophobic residues towards the membrane interface more favorably than other existing βArr1 complexes (6PWC, 6TKO), which is expected to promote membrane interaction (Lally et al., 2017). The inactive conformation of the finger loop was obtained from the inactive structure of βArr2 (PDB code: 3P2D) (Zhan et al., 2011).

To study spontaneous interaction with the plasma membrane, the structure of βArr2 was placed in proximity to the lipid bilayer. To mimic the experimental conditions as much as possible, we have taken into account the membrane composition of CHO cells (Symons et al., 2021) and incorporated the five most abundant membrane components into our simulation setup: 10% cholesterol, 38% palmitoyl-oleoyl-phosphatidylcholine, 28% dioleoyl-phosphatidylcholine, and 24% dioleoyl-phosphatidylethanolamine.

Membrane building and system solvation were done using the CHARMM-GUI server (Jo et al., 2008; Wu et al., 2014). The ionic strength of the system was kept at 0.15 M using NaCl ions.

Simulations were performed with the ACEMD3 package (Harvey et al., 2009). Parameters for system components were obtained from CHARMM36m (Huang et al., 2017) and CHARMM36 force fields (Best et al., 2012; Klauda et al., 2010). The simulation data are made available via the GPCRmd platform (Rodríguez-Espigares et al., 2020).

All the simulated systems were first relaxed during 50 ns of simulations under constant pressure and temperature (NPT) with a time step of 2 fs, with harmonic constraints applied to the protein backbone. The temperature was maintained at 310 K using the Langevin thermostat (Grest and Kremer, 1986) and pressure was kept at 1 bar using the Berendsen barostat (Eslami et al., 2008). The equilibration run was followed by production runs in conditions of constant volume and temperature (NVT) with a 4-fs time step. No constraints were applied during this stage. In all simulations, we used van der Waals and short-range electrostatic interactions with a cut-off of 9 Å and the particle mesh Ewald method (Darden et al., 1993) for long-range electrostatic interactions. The resulting simulation frames were analyzed using VMD (Humphrey et al., 1996) and tools available within, as well as in-house scripts.

To study spontaneous plasma membrane insertion of βArr2, 40 x production runs of 60 ns were carried out, starting from a conformation of βArr2 in the vicinity of the membrane. Afterwards, we carried out 4 x additional simulations, each amounting to 600 ns, starting from a C-edge anchored conformation of βArr2 observed in the previous simulations, in which we were able to observe penetration of the membrane by the finger loop.

To further study interaction of βArr2 with the membrane, we carried out 3 x 1 μs of production runs, starting from the fully membrane-anchored conformation obtained in spontaneous association experiments. To generate the βArr2 lipid-mutant, the involved modifications were introduced with the CHARMM-GUI server.

#### Metadynamics

Due to the high complexity and huge degrees of freedom of the arrestin–membrane ensemble, the simulation time required to obtain a converged energetic estimate of the membrane (un)binding process would exceed our computational capabilities. To overcome this limitation, we simulated the membrane (un)binding of the C-domain of βArr2 (P176-P347) alone, with or without the investigated mutations. Even though these simulations do not include the whole arrestin–membrane ensemble, they provide valuable insights into the effects of mutations on the free energy associated with membrane (un)binding.

All well-tempered metadynamics simulations were performed using ACEMD3.3 with the plumed 2.6.1 plugin.

The collective variable (CV) used to bias the simulations was the distance (*z*_1_-*z*_2_) between the mean position of phosphorous atoms (*z*_1_ coordinate) in the lower leaflet of the membrane (cytoplasmic surface) and the geometric center of the Cα positions in the C-domain of βArr2 (*z*_2_ coordinate). The choice of this collective variable ensures sampling of only the βArr2 (un)binding event, while excluding the movements of βArr2 along the xy plane of the membrane. An upper wall for the CV was set at a distance of 7.5 nm with a force constant of *K* = 500 KJ·mol^-1^·nm^-2^, aiming to avoid the escape of the C-domain into the bulk solvent and, thus, helping the convergence of the metadynamics simulations and reducing the computational burden.

The starting structure for the metadynamics simulations had the C-domain bound to the membrane and was obtained by equilibrating the C-domain for 50 ns under NPT conditions. The same general parameters (time step, thermostat, etc.) of the unbiased MD simulations were used. Metadynamics simulations were performed using an initial bias height of 1 KJ·mol^-1^, a width of 0.5, a bias factor of 16 and the rate of Gaussian deposition set to 500 steps. We performed 2 independent well-tempered metadynamics simulations for both wild-type and mutant C-domain until they reached convergence.

To check for convergence, we performed a block analysis of the histogram of the CV, which showed (for both simulations) a hyperbolic-like graph, suggesting that the system had converged at 1.8 µs. To compute the free energy, we followed the protocol described by Bussi and Laio (Bussi and Laio, 2020) for well-tempered metadynamics, and we estimated the error on the free energy by using the previous block analysis.

#### Contact analysis

To study the strength and stability of βArr2–membrane interactions, we carried out 3 production runs of 1 μs each, starting from the conformation obtained in spontaneous association experiments with βArr2 anchored to the lipid bilayer.

To quantify the interactions, we monitored the distance from the membrane of reference amino acids in the finger loop (A61-D79), C-loop (A240-Q251) and C-edge (T188-H199, N224-E231, S329-V336), using the following rational switching function:

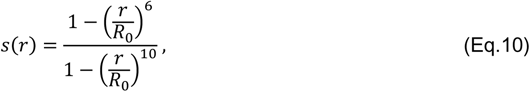

where *R*_0_ is 0.5 nm and r is the distance between the geometric center of the sidechain of each amino acid (Hα1 and Hα2 for glycine) and all the atoms composing the membrane. The main reason for using the geometric center of each sidechain instead of the position of the contained amino acids is to normalize for the differences in the number of sidechain atoms between amino acids.

The coordination number *N* for each amino acid is defined as the sum over all the atoms of the membrane in the simulation according to the following expression:

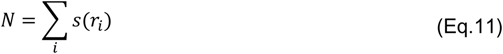

By using this approach, we can quantify the coordination number for each of the residues studied, which is directly related to the strength of the interaction between βArr2 and the lipid bilayer. The coordination number is calculated for each residue in each frame for the three accumulated microseconds to posteriorly calculate its mean value for the selected finger loop, C-loop and the C-edge residues. To simplify the comparison between wild-type and mutant βArr2, total coordination numbers for the finger loop, C-loop and C-edge were calculated by summing up the coordination numbers of each of the residues in each region.

#### Inter-domain rotation angle

To study the impact of membrane anchoring on the conformation of βArr2, we carried out 3 additional 1 μs runs with βArr2 in solution. The rotation angle between βArr2 N- and C-domains was computed as previously described (Dwivedi-Agnihotri et al., 2020). The utilized scripts were kindly provided by Naomi Latoracca (Latorraca et al., 2018). To evaluate the ability of Fab30 to recognize the observed active-like conformation of the fully membrane engaged βArr2, we compared the conformations obtained in MD simulations with that of the structure of βArr1 in complex with Fab30 and a fully phosphorylated 29-amino-acid C-terminal peptide derived from the vasopressin V_2_ receptor (PDB code: 4JQI) (Shukla et al., 2013).

#### Quantification and statistical analysis

Statistical analyses were performed using MATLAB (version 2018b). Differences between three or more groups were assessed by a non-parametric Kruskal Wallis test followed by unpaired two-tailed t-test with Bonferroni correction. Differences were considered significant for P < 0.05. Single-molecule data were analyzed by automated scripts with no user intervention during the analysis. Statistical parameters are reported in the figure legends.

## SUPPLEMENTAL INFORMATION

**Table S1.**
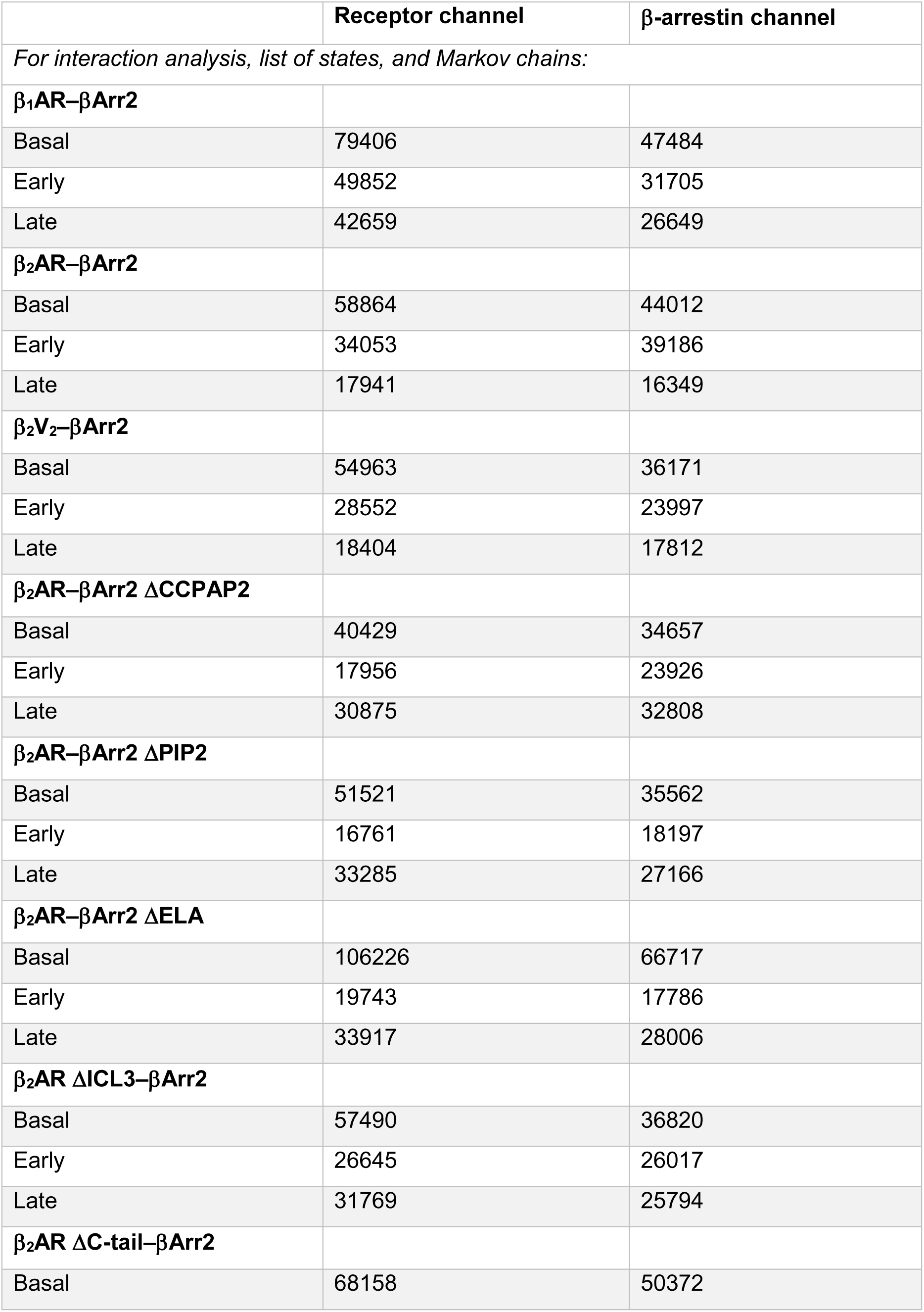
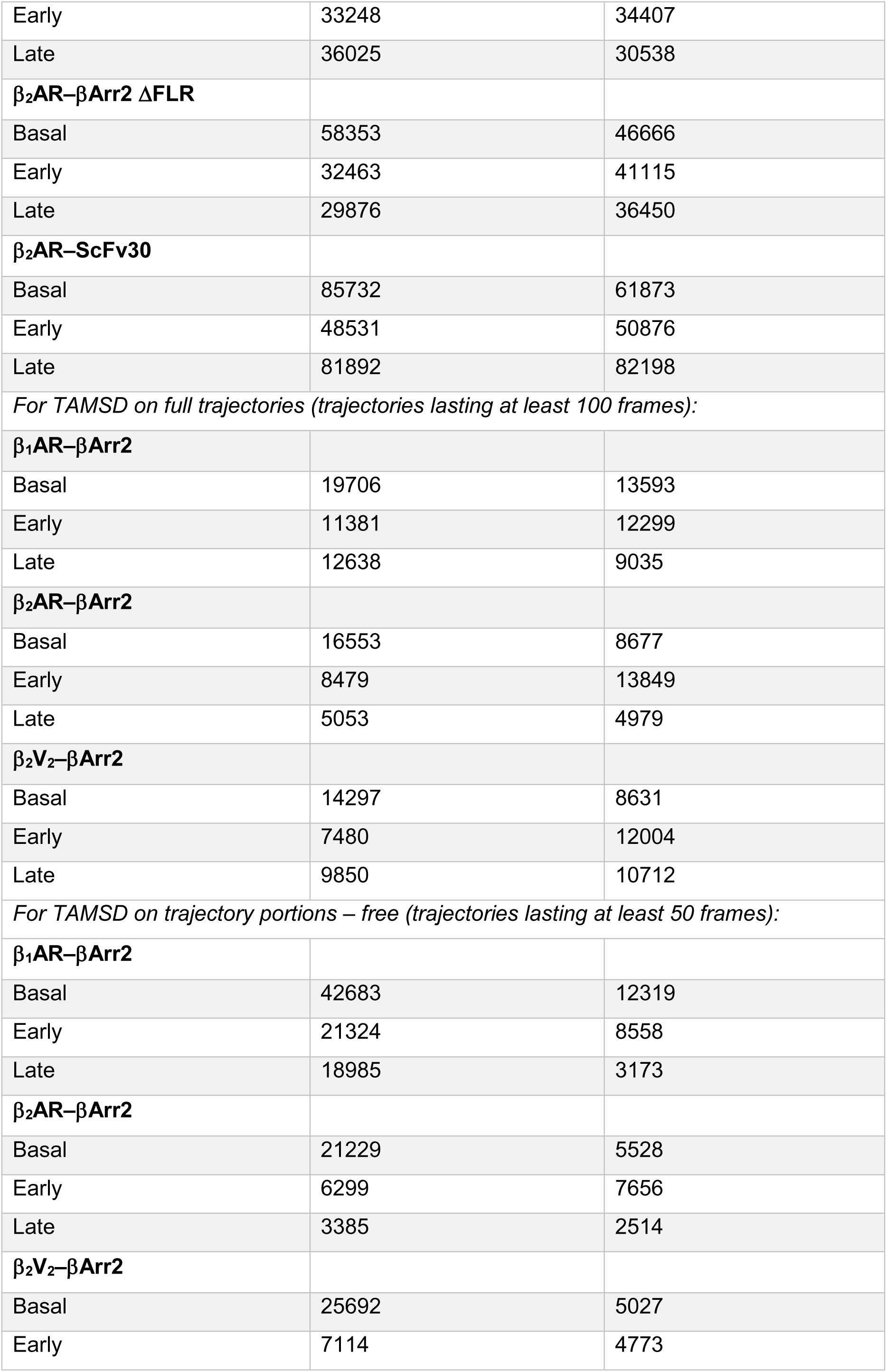
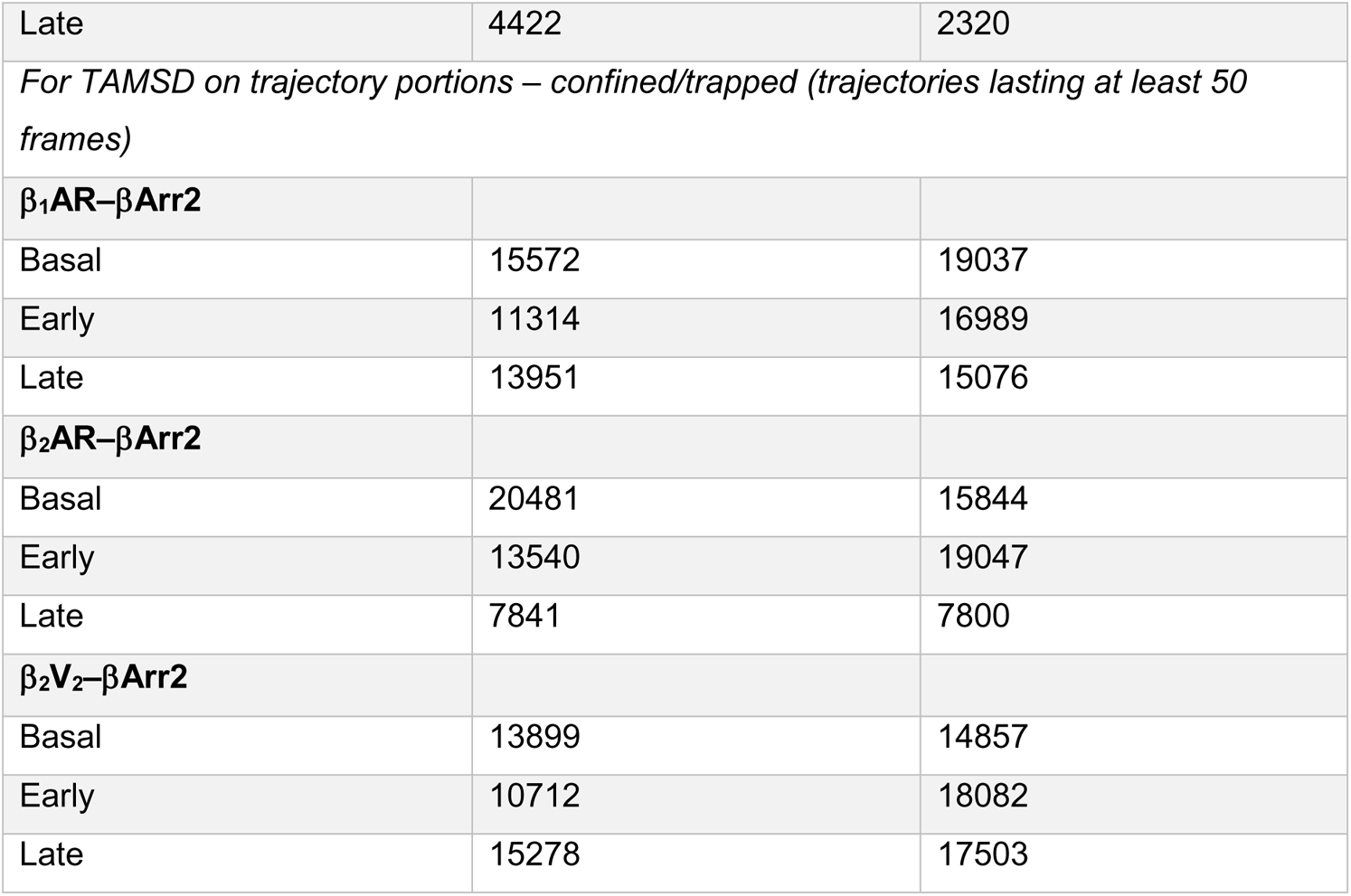
**Numbers of Analyzed Single-Molecule Trajectories for Each Condition**

**Table S2.**
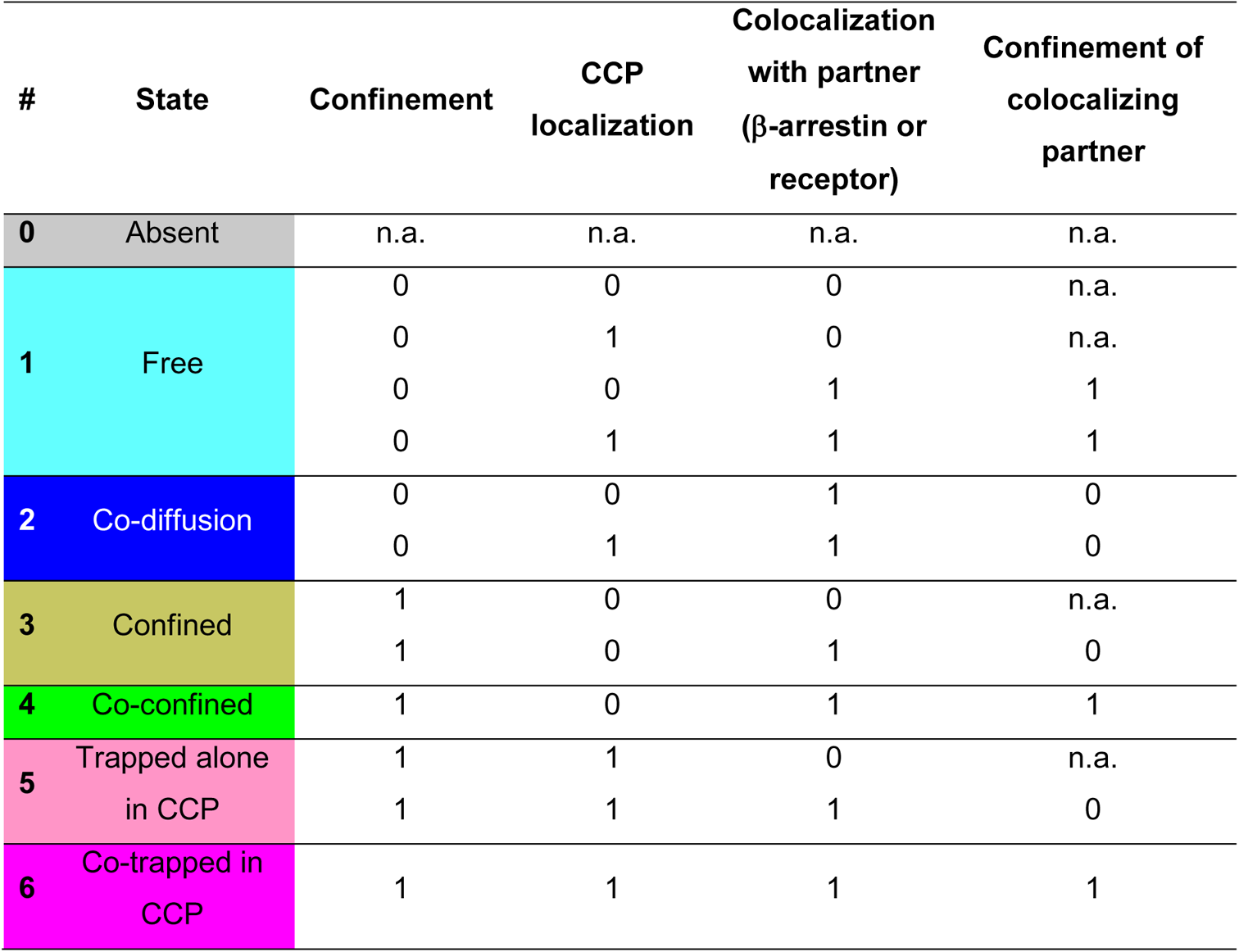
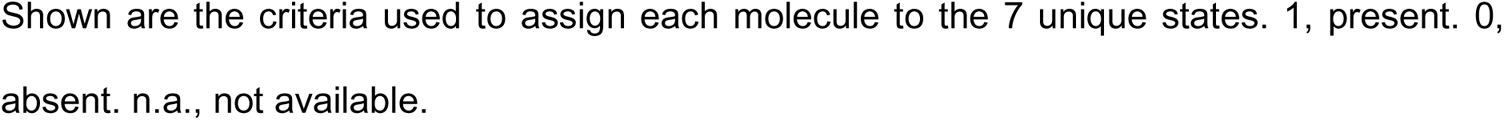
**States Used in Markov Chain Analyses**

**Table S3.** **Details of MD Simulations**

| Simulated transition | Replicates | Starting conformation | No. of times observed |
| --- | --- | --- | --- |
| Membrane penetration by C-edge (Transition 1) | x 40<br>[60 ns] | βArr2 not contacting membrane | 4 / 40 |
| Membrane penetration by finger loop (Transitions 2 & 3) | x 4<br>[200 ns] | βArr2 anchored via the C-edge | 1 / 4 |

## Supplemental Videos

**Video S1. Individual** β**_2_AR and** β**Arr2 Molecules Diffusing on the Plasma Membrane of a Living Cell**

Shown is a representative two-color TIRF image sequence of β_2_AR (green) and βArr2 (magenta) with trajectories overlaid. Frames were acquired every 33 ms. Playback, 30 frames/s.

**Video S2.** β**_2_AR and** β**Arr2 Single-Molecule Trajectories**

Shown are the same trajectories of Video S1 overlaid on the corresponding CCP binary mask. Frames were acquired every 33 ms. Playback, 15 frames/s.

